# CDK1 and PLK1 cooperate to regulate BICD2, dynein activation and recruitment to the nucleus as well as centrosome separation in G2/M

**DOI:** 10.1101/2021.11.04.467245

**Authors:** Núria Gallisà-Suñé, Paula Sànchez-Fernàndez-de-Landa, Fabian Zimmermann, Marina Serna, Joel Paz, Laura Regué, Oscar Llorca, Jens Lüders, Joan Roig

## Abstract

The activity of dynein is regulated by a number of adaptors that mediate its interaction with dynactin, effectively activating the motor complex while also connecting it to different cargos. The regulation of adaptors is consequently central to dynein physiology, but remains largely unexplored. We now describe that one of the best-known dynein adaptors, BICD2, is effectively activated through phosphorylation. In G2 phosphorylation of BICD2 by CDK1 promotes its interaction with PLK1. In turn, PLK1 phosphorylation of a single residue in the N-terminus of BICD2 results in a conformational change that facilitates interaction with dynein and dynactin, allowing the formation of active motor complexes. BICD2 phosphorylation is central for dynein recruitment to the nuclear envelope, centrosome tethering to the nucleus and centrosome separation in G2/M. This work reveals adaptor activation through phosphorylation as crucial for the spatiotemporal regulation of dynein activity.

## Introduction

Cytoplasmic dynein-1 (referred to herein simply as “dynein”) is a highly conserved microtubule motor that has a wide array of functions both in dividing and non-dividing cells (Reck-Peterson et al., 2018; Canty et al., 2021). As the primary minus-end-directed motor in animal cells, dynein participates in the transport of different types of vesicles, proteins and mRNA particles. Viruses can be similarly transported by the motor during infection. In addition, as a result of its ability to exert force on cellular structures, dynein contributes to the positioning of organelles such as centrosomes, the Golgi apparatus and the nucleus (Tanenbaum et al., 2011; Yadav and Linstedt, 2011; Agircan et al., 2014), and has an important role during mitotic spindle formation (Raaijmakers and Medema, 2014).

Dynein is an oligomer of two copies each of six different proteins: dynein heavy chain (DHC, containing both a motor and a microtubule-binding domain), an intermediate chain (DIC), a light intermediate chain (DLIC), and three different light chains (DLC). Their structural organization allows the resulting complex to move and exert forces along microtubules (Carter et al., 2016; Canty et al., 2021). Dynein on its own is in an autoinhibited conformation. Its mobility, processivity and ability to exert force are greatly increased by its interaction with dynactin plus one of different adaptors. Dynactin is a multiprotein complex that contains an actin-like filament and a shoulder region that interacts with microtubules through the extended arm of its p150^Glued^ subunit (hereinafter referred to as p150). The dynein adaptors are non-related coiled-coil rich proteins that facilitate the interaction between one or two dynein complexes with dynactin, effectively acting as activators of the motor complex. Adaptors do so by providing a ∼300 residue long region that is positioned in an extended conformation along the dynactin filament while also interacting with dynein, thus organizing the motor complex in a manner that favors mobility and processivity. Dynein adaptors are additionally in charge of connecting the motor complex with different cellular cargos, resulting in their transport along microtubules (Reck-Peterson et al., 2018; Olenick and Holzbaur, 2019).

One of the most studied dynein adaptors is BICD2. It belongs to a family of proteins related to *Drosophila* BICD, originally described as a polarity factor and subsequently found to work alongside dynein and different partners to localize a variety of cellular components in a microtubule-dependent manner (see (Claussen and Suter, 2005) for a review). In humans, the BICD family contains 4 members, BICD1, BICD2, BICDL1 and BICDL2 (also known as BICDR1/2), all of which have been described as dynein adaptors in different cellular contexts (Hoogenraad and Akhmanova, 2016). BICD proteins contain long coiled coil regions and function as dimers. They are able to bind dynein and dynactin through an extended rod-shaped region in their N-terminus, while interacting with different cargos through their C-terminus. BICD2 is a ∼94 kDa protein that similarly to the closely related BICD1 is composed of three coiled coil regions (named CC1-CC3) separated by two and flexible linkers predicted to be unstructured. BICD2 interacts with dynein and dynactin through its most N-terminal CC1 region and part of the central CC2 region (Hoogenraad et al., 2003; Splinter et al., 2012). This segment has a rod-shaped structure and is sandwiched between dynein and dynactin (Urnavicius et al., 2015). The CC1 contains a motif aptly named CC1 box, which is conserved across the BICD family, and that is crucial for the interaction with dynein DLIC. Additionally the CC2 contains a so-called Spindly motif, which interacts with the pointed end of the dynactin filament (Gama et al., 2017; Lee et al., 2018). When separated from the rest of the protein, an N-terminal fragment of BICD2 containing the mentioned elements is able to induce minus-end-directed mobility of dynein/dynactin complexes (Hoogenraad et al., 2001, 2003; Splinter et al., 2012) by activating and making them processive (McKenney et al., 2014; Schlager et al., 2014).

Importantly, BICD2 CC3 interacts with the CC1 region, reducing the dynein-binding activity of the N-terminus, thus effectively acting as an auto-inhibitory domain (Hoogenraad et al., 2001, 2003). The same region also interacts with Rab6, a small GTPase involved in the control of membrane traffic, linking the adaptor to the regulation of Golgi structure and vesicle transport in the exocytotic pathway (Matanis et al., 2002; Grigoriev et al., 2007). The current model for the regulation of BICD2 (and possibly other BICD family members) suggests that binding of Rab6 to the C-terminus releases the N-terminal region from its embrace, “opening” the molecule and favoring the formation of active complexes with dynein and dynactin (Hoogenraad et al., 2001; Matanis et al., 2002; Hoogenraad and Akhmanova, 2016; Huynh and Vale, 2017). Rab6 is thought to be the major cargo of BICD2 during G1 and S (and in non-dividing cells), and accordingly the adaptor localizes at the Golgi during these phases of the cell cycle. In early G2, however, BICD2 accumulates at the nuclear envelope where it recruits dynein and dynactin. This is the result of BICD2 switching cargos and interacting with the nucleoporin RanBP2/NUP358 (Splinter et al., 2010). A complementary, non-redundant dynein-recruitment pathway exist during late the G2 and early M phases, mediated by Nup133, CENPF and NDE1/NDEL1 (Bolhy et al., 2011). Dynein at the nuclear envelope plays a major role in tethering the centrosomes and the nucleus, contributing both to early centrosome separation and proper spindle assembly (Gönczy et al., 1999; Robinson et al., 1999; Splinter et al., 2010; Bolhy et al., 2011; Raaijmakers et al., 2012), as well as to nuclear envelope breakdown (Salina et al., 2002). Interestingly, in neural progenitors it also drives apical nuclear migration (Hu et al., 2013). Dynein recruitment is controlled in neural and non-neural cell types by the cyclin-dependent kinase CDK1, which phosphorylates RanBP2 to induce its binding to BICD2, as well as CENP-F and NDE1 thus controlling the Nup133-based pathway (Baffet et al., 2015; Wynne and Vallee, 2018). Whether other inputs besides CDK1 control dynein recruitment to the nuclear envelope is not known; published data suggest that the Polo-family kinase PLK1 may also have a role in this regulation, although the molecular details of this are not known (Baffet et al., 2015).

Here we present a new regulatory mechanism controlling the recruitment of dynein to the nuclear envelope that is based on the sequential phosphorylation of the dynein adaptor BICD2 by the mitotic kinases CDK1 and PLK1. Our results show that BICD2 interacts with PLK1 through the kinase polo-box domain (PBD), and that this interaction depends on the phosphorylation of BICD2 by CDK1. In turn, PLK1 is able to phosphorylate Ser102 in the N-terminus of BICD2. We show that modification of Ser102 alters the conformation of BICD2, interfering with the interaction between BICD2 N- and C-terminal regions. Importantly, Ser102 modification increases the ability of BICD2 to bind and activate dynein. We highlight the physiological relevance of our observations by showing that the phosphorylation of BICD2 by CDK1 and PLK1 is necessary for dynein recruitment to the nuclear envelope and proper nucleus-centrosome tethering in G2, as well as for subsequent centrosome separation in early mitosis. In summary, we identify adaptor phosphorylation as a novel mechanism to regulate dynein localization and activation.

## Results

### PLK1 PBD interacts with BICD2 in a CDK phosphorylation-dependent manner

Several high throughput studies indicate that BICD2 is phosphorylated *in vivo* at multiple residues (see https://www.phosphosite.org/ and studies cited therein). We wondered whether some of these posttranslational modifications could be modulating BICD2 function, and sought to identify the kinase responsible for them. A good candidate was PLK1, a protein kinase active during G2 and M and with multiple functions during these phases of the cell cycle (Bruinsma et al., 2012; Zitouni et al., 2014). PLK1 had been previously suggested to interact with BICD2 by various assays including yeast two hybrid (Mirouse et al., 2006), affinity purification coupled to mass spectrometry (MS) (Hutchins et al., 2010) and proximity labeling-MS (Redwine et al., 2017)). Moreover, some of the BICD2 phosphosites found *in vivo* conform to a putative PLK1-phosphorylation motif (see below). We thus focused our efforts on this protein kinase. To further confirm that the two proteins could indeed form a complex in mammalian cells, we immunoprecipitated endogenous BICD2 from exponentially-growing and mitotic HeLa cells and probed the precipitates with anti-PLK1 antibodies (Figure 1A). Indeed, PLK1 was specifically detected in BICD2 immunoprecipitates and the detected interaction was substantially increased in mitosis, when PLK1 is active. Interestingly in this phase of the cell cycle BICD2 showed a higher apparent molecular weight after electrophoresis (see also Figure 1C), thus suggesting that it was phosphorylated. Additional immunoprecipitations using recombinant fragments of BICD2 indicated that PLK1 preferentially interacted with the region containing the first two coiled-coil regions of the adaptor (BICD2 [1-575]; note that unless otherwise indicated, BICD2 residue number in this work refer always to the mouse sequence) (Figure 1B).

**Figure 1.**
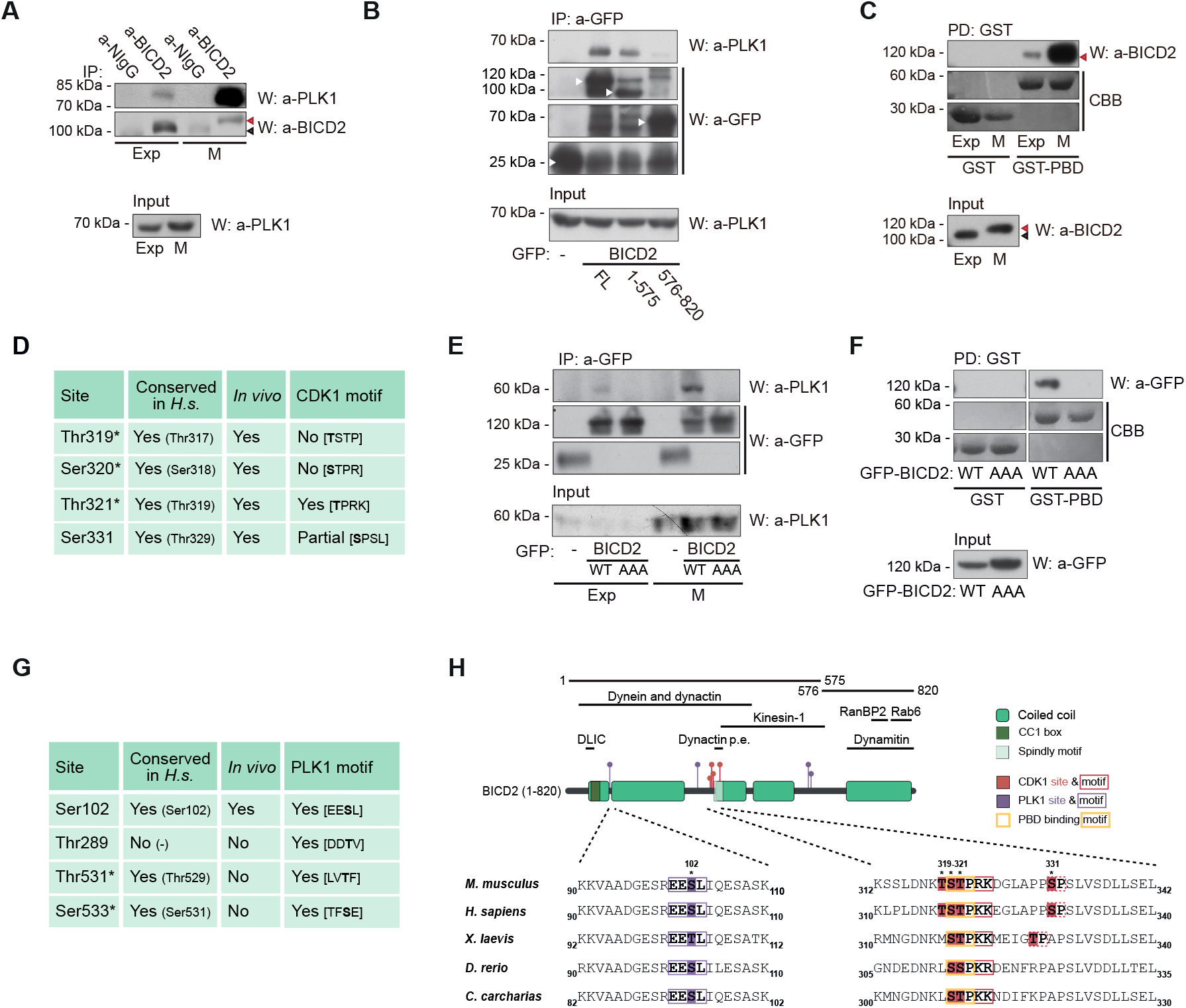
BICD2 interacts with PLK1 through its Polo-Box Domain and is a PLK1 and CDK1 substrate. **(A)** Coimmunoprecipitation of endogenous PLK1 and BICD2 in exponentially growing (*Exp*) and nocodazole-arrested mitotic (*M*) HeLa cell extracts. anti-BICD2 or normal IgG (*NIgG*) immunoprecipitates (*IP*) were analyzed by western blot (*W*) using either anti-PLK1 or anti-BICD2 antibodies. Note BICD2 high apparent molecular weight in mitotic cells, arrowheads. PLK1 in the corresponding extracts is shown in the lower panel. **(B)** PLK1 interacts with the N-terminal region of BICD2. Immunoprecipitates of the indicated recombinant GFP-fusion proteins (arrowheads) were analyzed by western blot using either anti-PLK1, to detect the endogenous kinase, or anti-GFP. PLK1 in the corresponding extracts is shown in the lower panel. **(C)** PLK1 interacts with BICD2 through its Polo-Box Domain (PBD). Extracts of exponentially growing (*Exp*) or nocodazole-arrested mitotic (*M*) HeLa cells were incubated with either bacterial expressed GST or GST-PBD (GST-PLK1 [354-603]) bound to GSH beads. Endogenous BICD2 was detected by western blot (*W*) and GST-fusion proteins by Coomassie Brilliant Blue staining (*CBB*). BICD2 in the corresponding extracts is shown in the lower panel. Note that in both pulldowns from exponential and mitotic extracts the apparent molecular weight of BICD2 interacting with PLK1-PBD corresponds to the fastest migrating form (red arrowhead). **(D)** Sites phosphorylated *in vitro* in BICD2 [272-540] by CDK1, as identified by LC-MS/MS (see also Supplementary Figure S1A). *Conserved in H.s.*, conservation in humans (the orthologous residue is indicated between parenthesis)*. In vivo,* sites previously determined to be phosphorylated *in vivo* in human cells (see https://www.phosphosite.org/). *CDK1 motif,* presence of a CDK1 consensus motif around the phosphorylation site ([S/T]PX[K/R] where X is any residue and S or T are the modified residues). Asterisks denote that a single phosphorylation site was detected that could not be assigned unequivocally among the three residues. **(E)** and **(F)** The interaction of BICD2 and PLK1 depends on the residues phosphorylated by CDK1. (E) anti-GFP immunoprecipitates from extracts of exponentially growing (*Exp*) or nocodazole-arrested mitotic (*M*) HeLa cells expressing either GFP, GFP-BICD2 wild type (*WT*) or GFP-BICD2 [T319A, S320A, T321A] (*GFP-BICD2 AAA*). Endogenous PLK1 and GFP are detected by western blot. PLK1 in the corresponding extracts is shown in the lower panel. (F) Extracts of HeLa cells expressing GFP-BICD2 wild type or GFP-BICD2 AAA were incubated with GST or GST-PBD bound to GSH beads. GFP was detected by western blot (*W*) and GST-fusion proteins by Coomassie Brilliant Blue (*CBB*) staining. **(G)** Sites phosphorylated *in vitro* in BICD2 [1-271] and BICD2 [272-540] by PLK1, as identified by LC-MS/MS (see also Supplementary Figure S2A). *Conserved in H.s.*, conservation in humans (the orthologous residue is indicated between parenthesis). *In vivo*, sites previously determined to be phosphorylated in vivo in human cells (see https://www.phosphosite.org/). *PLK1 motif*, presence of a PLK1 consensus motif around the phosphorylation site ([D/E]X[S/T]Φ, where X is any aminoacid and Φ is an hydrophobic residue). Asterisks denote that a single phosphorylation site was detected that could not be assigned unequivocally among the two residues. **(H)** *Top*, schematic representation of BICD2, noting regions interacting with different proteins or protein complexes, plus the different fragments and residues mentioned in this figure (*Dynactin p.e., dynactin pointed end*). Residue number corresponds to mouse BICD2. Coiled coil regions have been predicted using paircoil2 (McDonnell et al., 2006). *Bottom*, sequence alignments of residues surrounding Ser102, Thr319, Ser320, Thr321 and Ser331 in different BICD2 vertebrate orthologues are shown: mouse (Q921C5), human (Q8TD16), *Xenopus laevis* (Q5FWL8), *Dario rerio* (X1WDT9) and *Carcharodon carcharias*, a cartilaginous fish (XP_041047193). See also Supplementary Figure S2B for similar alignments with BICD family members.

PLK1 interacts with its substrates in a phosphospecific manner through its polo-box domain (PBD) (Elia et al., 2003a). We thus expressed GST-PLK1 PBD (GST-PLK1 [354-603]) in bacteria and determined whether it was able to interact with BICD2 from cell extracts. We indeed observed that GST-PLK1 PBD specifically interacted with BICD2 (Figure 1C). Moreover, and in agreement with the results obtained with full length PLK1, the amount of BICD2 associated with PLK1 PBD was significantly higher when mitotic extracts were used. Additionally, we noted the PBD almost exclusively interacted with the form of BICD2 that showed a high apparent molecular weight, prevalent in mitotic cells. Altogether our results indicate that PLK1 interacts through its PBD domain with modified BICD2, though a region in the N-terminal part of the adaptor. This is in agreement with results obtained by Mirouse and collaborators using the yeast two-hybrid technique (Mirouse et al., 2006).

The PBD is a phosphoserine/phosphothreonine binding domain that depends on a priming phosphorylation step for its interaction with other proteins, frequently carried out by proline-directed kinases such as the cyclin-dependent kinases (CDKs) (Elia et al., 2003a). Our results suggest that BICD2 may preferentially interact with PLK1 in G2/M and thus we inquired whether CDK1, in charge of controlling progression through these phases of the cell cycle together with CDK2, could modify BICD2. For this, we sought to express three different fragments of BICD2 in bacteria containing each one of the coiled-coil regions of the adaptor. We succeeded in purifying as GST-fusions the two most N-terminal fragments (GST-BICD2 [1-271] and GST-BICD2 [272-541]), comprising most of the region shown to bind PLK1 in Figure 1B. Upon incubation with CDK1/cyclin B plus ATP/Mg^2+^, GST-BICD2 [272-541] became readily phosphorylated, in contrast to GST-BICD2 [1-271] that was only slightly modified (Supplementary Figure S1A). Proteolytic digestion followed by liquid chromatography-tandem MS (LC-MS/MS) analysis identified two phosphorylated peptides in GST-BICD2 [272-541]. One of the modifications was unequivocally assigned to a single residue, Ser331, while the other could not be assigned among three contiguous residues (_319_TST_321_) (Figures 1D and 1H). All of the CDK1 putative sites cluster in close proximity and have been previously shown to be phosphorylated in humans *in vivo* (see https://www.phosphosite.org/). Mutation of Thr319, Ser320 and Thr321 to non-phosphorylatable alanines (GST-BICD2 [272-541; T319A, S320A, T321A]) highly impaired phosphorylation by CDK1/cyclin B, thus suggesting that at least one of the three residues is the major target of the kinase in the N-terminal region of BICD2 (Supplementary Figure S1B). Thr319, Ser320 and Thr321 and its surrounding residues are mostly conserved in mammals and in other vertebrates (Figure 1H). Importantly, they contain a canonical CDK phosphorylation motif ([S/T]PX[K/R], where X is any amino acid; for example (Songyang et al., 1994)) and, when modified, conform to a PBD-interacting motif (S[pS/pT]PX, where pS and pT are phosphoserine and phosphothreonine, respectively; (Elia et al., 2003a)). Our results thus support the hypothesis that CDK1 phosphorylation of these residues could be regulating PLK1 binding to BICD2.

To test this, we next expressed both wild type BICD2 and a form of BICD2 that cannot be phosphorylated at _319_TST_321_ (BICD2 [T319A, S320A, T321A], henceforth BICD2 AAA) in mammalian cells and determined whether mutation of these residues would interfere with PLK1 binding to BICD2. Indeed, we observed that the GFP-BICD2 AAA mutant did not interact with PLK1 in neither exponentially-growing or mitotic cells (Figure 1E). Moreover GFP-BICD2 AAA was not pulled down from cell extracts by GST-PLK1 PBD, in contrast to its wild type counterpart (Figure 1F). We interpret our data as a confirmation that CDK1 phosphorylation of one or more residues in BICD2 Thr319-Thr321 results in the interaction of BICD2 with PLK1 through the PLK1 PBD.

### PLK1 phosphorylates Ser102 in the N-terminal dynein-interacting region of BICD2

As an initial step to study whether PLK1 may have a role regulating BICD2 function, we sought to determine whether the adaptor is a PLK1 substrate. For this we incubated GST-BICD2 [1-271] and GST-BICD2 [272-541] with PLK1 plus ATP/Mg^2+^ (Supplementary Figure S2A). Both polypeptides were phosphorylated by PLK1, although GST-BICD2 [1-271] was a better substrate, incorporating rapidly up to three times more phosphate than GST-BICD2 [272-541] upon incubation with PLK1. Trypsin digestion followed by LC-MS/MS analysis identified three BICD2 peptides phosphorylated by PLK1. Two of the modifications were unequivocally assigned to Ser102 and Thr289, while the third could not be assigned among two nearby residues, Thr531 and Ser533 (Figures 1G and 1H). Of the identified sites, only Ser102 was located in GST-BICD2 [1-271] suggesting that the modification of this residue accounted for most of the phosphate incorporated in the adaptor *in vitro*. BICD2 Ser102 is conserved in jawed vertebrates (*Gnathostomata*), including tetrapods as well as bony and cartilaginous fish (Figure 1H), and is surrounded by a sequence that conforms to a canonical PLK1 phosphorylation site ([D/E]X[S/T]Φ, where X is any aminoacid and Φ is an hydrophobic residue (Nakajima et al., 2003; Alexander et al., 2011). The residue and surrounding aminoacids are conserved as well in BICD1 (Supplementary Figure S2B). Additionally, among the phosphosites identified in our study Ser102 is the only one that has been detected *in vivo* in human cells. We thus focused our study on the functional role of the phosphorylation of this BICD2 residue.

### Modification of BICD2 Ser102 favors dynein binding and activation

Ser102 is located at the N-terminal region of BICD2, close to the dynein-interacting CC1 box (see Figure 1H). We reasoned that phosphorylation could regulate the ability of this region to form a complex with the dynein motor complex. To investigate this, we mutated Ser102 to non-phosphorylatable (BICD2 S102A) and phosphomimetic (BICD2 S102D) residues. When the different recombinant BICD2 forms were expressed in cells, we observed that the amount of dynein (detected using anti-DIC antibodies) and dynactin (detected with anti-p150 antibodies) immunoprecipitating with wild type GFP-BICD2 and GFP-BICD2 S102A was low and in some experiments barely detectable. GFP-BICD2 S102A tended to coimmunoprecipitate less dynein than the wild type form (Figure 2A). This is consistent with the suggestion that full length BICD2 exists in a “closed” conformation that precludes it from binding dynein/dynactin. GFP-BICD2 [1-575], lacking the cargo-binding regulatory C-terminal region, readily precipitated both dynein and dynactin as expected (Hoogenraad et al., 2003; Splinter et al., 2012). Importantly, the phosphomimetic BICD2 S102D consistently precipitated amounts of both DIC and p150 that were comparable to those associated with BICD2 [1-575], and that were significantly larger than those associated with wild type or S102A BICD2. Thus, the modification of Ser102 favors the interaction of full length BICD2 with dynein and dynactin. This was confirmed by experiments in which DIC was immunoprecipitated from cells expressing different forms of recombinant BICD2 (Figure 2B). In DIC immunoprecipitates, wild type and S102A GFP-BICD2 forms were below detection limits, whereas GFP-BICD2 S102D was readily detected. p150 was detected in all cases, possibly resulting from the presence in the cells of dynein/dynactin complexes formed with other adaptors, independently of BICD2.

**Figure 2.**
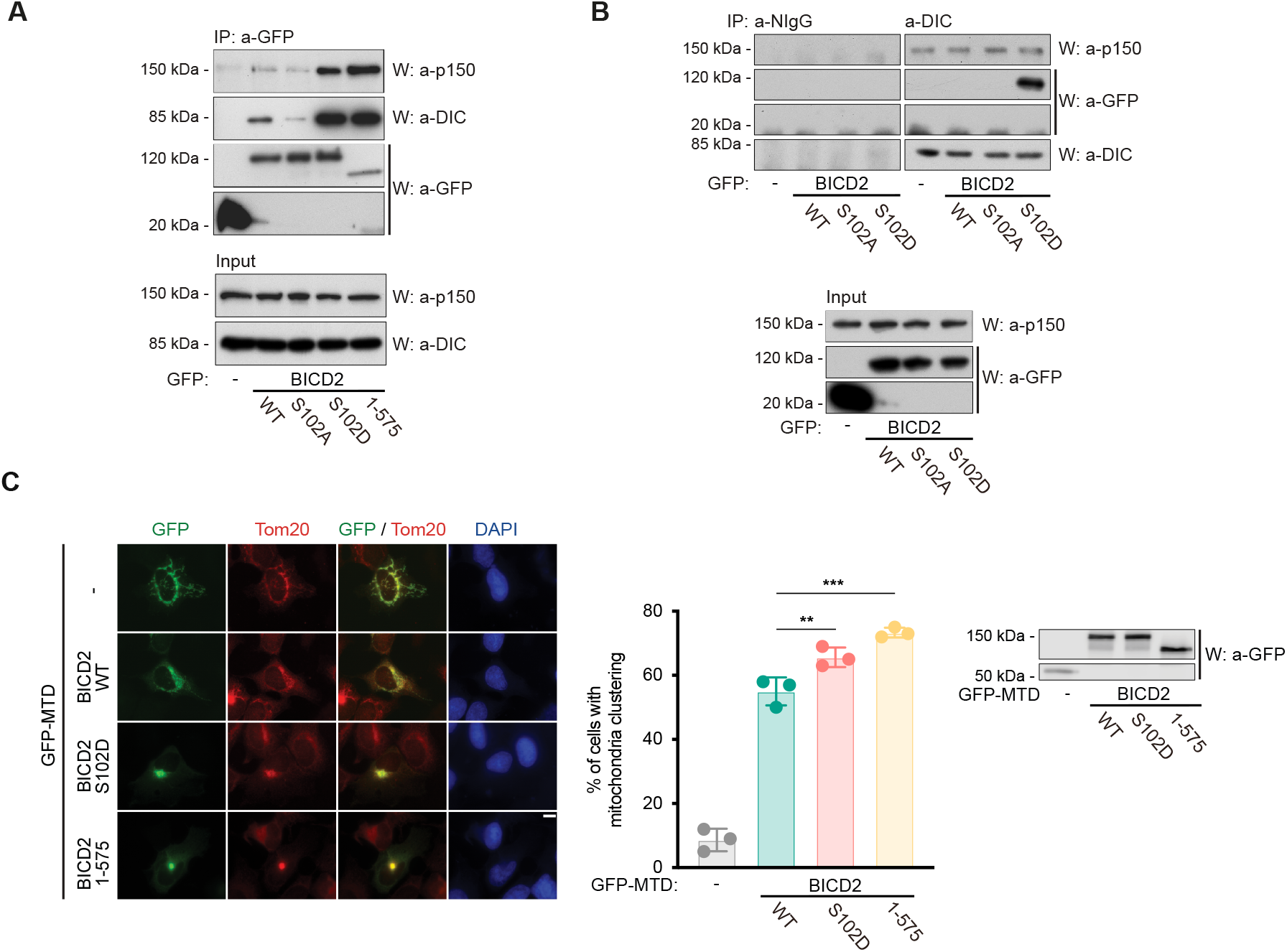
Modification of BICD2 at Ser102 is able to control dynein binding and mobility. **(A) and (B)** A phosphomimetic mutation in BICD2 Ser102 (BICD2 S102D) results in increased dynein and dynactin binding to BICD2. (A) The indicated GFP-fusion proteins were immunoprecipitated from HeLa cells and analyzed by western blot to detect dynein and dynactin (using respectively anti-dynein intermediate chain, DIC, and anti-p150 antibodies), plus GFP. DIC and p150 levels in the corresponding extracts are shown in the lower panel. (B) Normal IgG (NIgG) and anti-DIC immunoprecipitates from extracts expressing the indicated GFP-fusion proteins proved by western blot with the indicated antibodies. Levels of GFP-fusion proteins plus p150 in the corresponding extracts are shown in the lower panel. **(C)** A phosphomimetic mutation in BICD2 Ser102 (BICD2 S102D) results in increased dynein mobility towards the centrosomes as detected by mitochondria relocalization and clustering. Different GFP-fusion proteins were targeted to mitochondria through a mitochondria-targeting sequence domain (MTD) in U2OS cells. Representative immunofluorescence images are shown for each polypeptide, stained for GFP, Tom20, as a mitochondrial marker, and DAPI. Note that the S102D mutation favors clustering in a similar manner to the constitutively active BICD2 N-terminus (BICD2 1-575). Scale bar, 10 µm. See also Supplementary Figure S3A, showing that clustering happens around the centrosomes. Quantification of the percentage of cells with clustered mitochondria for each different GFP-fusion form is shown (n=3 biological replicates, 50 cells each; individual replicate means plus mean of replicates ± SD are shown; statistical significance analyzed using a Chi square test). Expression levels of the different polypeptides as detected by western blot are shown.

The interaction of dynein and dynactin with BICD2 has been shown to lead to the formation of a highly processive minus-end-directed motor complex (McKenney et al., 2014; Schlager et al., 2014). Therefore, we tested whether the modification of BICD2 Ser102 increased dynein mobility *in vivo*. For this, we used the ability of recombinant BICD2 to affect the localization of different organelles when artificially tethered to them. We fused different BICD2 forms to the mitochondria-targeting domain (MTD) of a splice variant of *Drosophila* centrosomin (Chen et al., 2017). When expressed in cells, the MTD fusion proteins co-localized with mitochondria as detected with the Tom20 mitochondrial marker. This allowed us to assess dynein activity by observing mitochondria clustering around the centrosome, where microtubules minus-ends converge. As expected from previous observations (Hoogenraad et al., 2003), expression of GFP-BICD2 [1-575]-MTD resulted in a dramatic clustering of the mitochondria in most cells (70%, Figure 2C). Clustering of mitochondria was around centrosomes as expected (see Supplementary Figure S3A). Wild type BICD2 (GFP-BICD2 WT-MTD) had a smaller effect, although surprisingly mitochondria clustering was still detected in about half of the cells (∼55%), suggesting that in the conditions used wild type GFP-BICD2-MTD was able to form active dynein complexes. Importantly, GFP-BICD2 S102D-MTD was significantly more efficient than wild type BICD2 in inducing the clustering of mitochondria (∼65% of cells with clustered mitochondria), thus supporting the hypothesis that the modification of Ser102 favors the formation of an active dynein motor complex.

### Modification of Ser102 induces conformational changes in BICD2

We reasoned that the modification of Ser102 could regulate the interaction of BICD2 with dynein by increasing its affinity for the motor complex, directly affecting BICD2 overall conformation (i.e. resulting in its “opening”) or both. To test the first possibility, we mutated Ser102 in BICD2 [1-575], devoid of regulation by the C-terminal part of the molecule. We observed that GFP-BICD2 [1-575; S102D] interacted with similar amounts of dynein as GFP-BICD2 [1-575] WT (or S102A) (Supplementary Figure S3B). Moreover, when expressed as GFP-MTD fusion proteins, both wild type and S102D forms of BICD2 [1-575] induced the clustering of mitochondria to an almost identical extent (Supplementary Figure S3C). This strongly suggests that the modification of Ser102 has an effect only in the context of the full-length molecule and does not directly increase the intrinsic affinity of the N-terminal part of BICD2 for dynein -and in this manner its ability to form an active motor complex.

We therefore sought to determine whether the modification of Ser102 could influence the molecular conformation of BICD2. For this we expressed TwinStrep-BICD2 WT and TwinStrep-BICD S102D in insect cells and purified both recombinant proteins to homogeneity. The two BICD2 forms showed almost identical profiles by size-exclusion chromatography combined with multiple angle light scattering (SEC-MALS), with a predicted MW of ∼190 kDa, thus showing that the mutation did not have an effect on dimerization and gross molecular size (data not shown). This was confirmed by dynamic light scattering (DLS), that indicated that both proteins had a similar hydrodynamic radius of ∼8nm (not shown). Also, we did not detect significant differences between wild type and mutated BICD2 when we measured their intrinsic fluoresce during a temperature ramp (not shown), suggesting that the mutation does not cause large structural rearrangements.

To directly assess the conformation of both forms we subsequently used negative stain electron microscopy (EM) (Figure 3A and B). We extracted several thousand individual molecule images of each sample that were subjected to image classification and 2D averaging. In the electron microscope wild type BICD2 appeared mostly as triangularly shaped molecules (68% of the total selected particles). These triangular images would be consistent with the shape of a BICD2 dimer where three rigid coiled coil regions are in a looped or closed conformation. Other less abundant images with a rod-like shape were also found (32% of the total selected particles). These could correspond to either a different conformation of BICD2 or a different view of the dimer when imaged from its side. Strikingly, when the S102D mutant was analyzed using similar methodologies, triangularly-shaped molecules were found to be less abundant (25% of the total selected particles) whereas rod-shaped molecules were now the major species (75% of the total selected particles). This strongly suggested that the mutation promotes changes in the conformation of BICD2. We cannot rule out completely that a major cause for the differences observed between the two proteins by EM are changes in view preference when the molecules interact with the support film. But even in this scenario, these changes would be a consequence of modifications in the structure of the protein caused by the phosphomimetic mutation.

**Figure 3.**
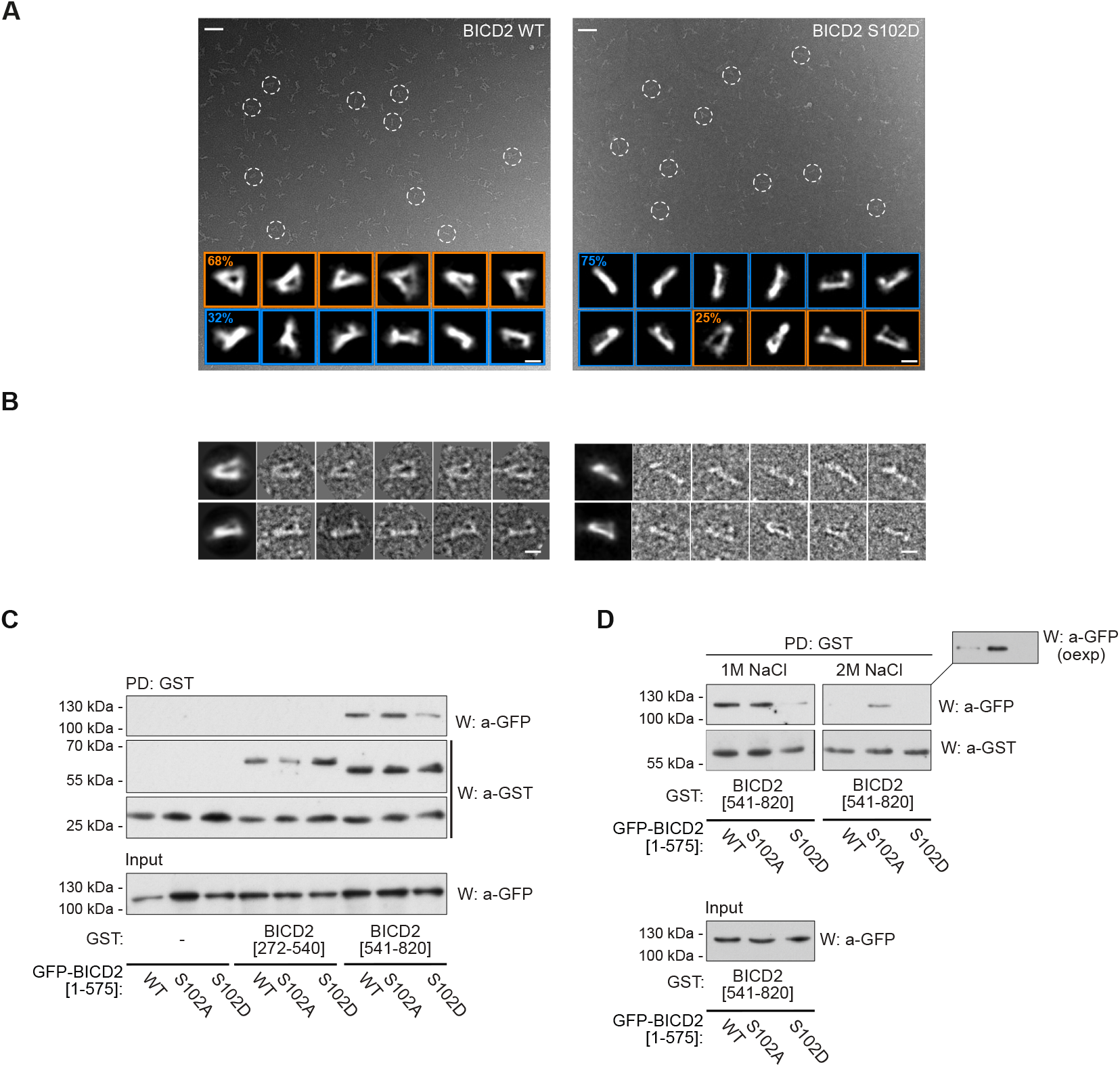
Modification of Ser102 alters the conformation of BICD2. **(A)** Representative electron micrographs obtained for wild type (*left*) and S102D (*right*) TwinStrep-BICD2, using negatively stained samples. Several BICD2 individual molecules are highlighted within white dashed circles. Representative 2D average images are displayed at the bottom of each micrograph. The percentage of particles with a triangular or a rod-like morphology is noted in each case (30,476 BICD2 wild type particles and 22,021 BICD2 S102D particles). Scale bars for micrographs and individual/average images represent 50 and 10 nm respectively. **(B)** Gallery of representative individual images of the wild type (left) and S102D (right) TwinStrep-BICD2 plus the corresponding average images. Scale bars, 10 nm. **(C)** Modification of Ser102 interferes with the interaction between BICD2 N- and C-terminal regions. HeLa cells were transfected with different forms of BICD2 N-terminus (BICD2 [1-575]) plus a fusion of GST and the C-terminal part of BICD2 (BICD2 [540-820]). GST alone or fused to an internal BICD2 region (GST-BICD2 [272-540]) were used as controls. GST pulldowns were analyzed by western blot (*W*) using either anti-GFP or anti-GST antibodies. GFP polypeptides in the corresponding extracts are shown in the lower panel. **(D)** A similar experiment in which 1M or 2M NaCl was added to the pulldown washing buffer (*oexp*, film overexposure).

After our EM observations, we sought to determine whether the modification of Ser102 could change the conformation of BICD2 by interfering with the interaction between its N-terminal and C-terminal parts. For this a GST-tagged fragment of BICD2 comprising the C-terminal CC3 region (GST-BICD2 [541-820]) was coexpressed with different GFP-tagged forms of GFP-BICD2 [1-575], containing both the CC1 and CC2 regions (see figure 1H). GST alone and GST-BICD2 [272-540] (containing the CC2 region) were used as controls. As expected from previous two-hybrid data (Hoogenraad et al., 2001), we observed that GST-BICD2 [541-820] (but not GST or GST-BICD2 [272-540]) was able to specifically pull down GFP-BICD2 [1-575] from cell extracts (Figure 3C). We interpret this as confirmation that the N-terminal and C-terminal region of the adaptor can interact in cells. Strikingly, the phosphomimetic form of the N-terminus GFP-BICD2 [1-575; S102D] but not the S102A form showed an impaired ability to interact with BICD2 C-terminus. This effect was enhanced when high salt concentrations were used during washes of the pulldowns (Figure 3D).

Altogether, we conclude that the phosphorylation of Ser102 directly interferes with the intramolecular interaction between the N-terminus and the C-terminus within the dimeric BICD2 molecule, resulting in a change in its conformation. The modified conformation, reflected as rod-like shapes in the EM images, is more prone to bind dynein and dynactin resulting in the formation of an active motor complex between them and the adaptor.

### Dynein binding to BICD2 in G2/M depends on PLK1 activity

Our results suggested that by phosphorylating BICD2 at Ser102, PLK1 may be able to regulate the formation of a functional complex between BICD2, dynein and dynactin in G2/M, when the kinase is active.

To explore this we determined the endogenous levels of dynein interacting with BICD2 in cells progressing unimpaired through G2, cells arrested in G2 after CDK1 inhibition with RO-3306 (Vassilev et al., 2006), and G2 cells in which both CDK1 and PLK1 where inhibited through the use of RO-3306 plus the PLK1 inhibitor BI 2536 (Steegmaier et al., 2007). Unfortunately we were not able to reliably detect endogenous dynactin (p150) in our immunoprecipitates of endogenous BICD2. Figure 4A shows that dynein readily coimmunoprecipitated with BICD2 in G2, with levels that were equal or slightly higher than those observed in exponentially growing or G1/S-arrested cells. G2 cells devoid of CDK1 activity showed diminished but still sizable interaction. Importantly, acute treatment (2h) with BI 2536 of G2-arrested cells totally abrogated the interaction between BICD2 and dynein. Our results thus show that the interaction in G2 is only partially dependent on the activity of CDK1, suggesting that another kinase such as CDK2 may be able to prime PLK1 binding to BICD2, or that PLK1 levels in cells arrested with RO-3306-arrested for several hours are enough to support BICD2 phosphorylation independently of the interaction. Importantly though, they also show that the interaction between BICD2 and dynein completely depends on the activity of PLK1 in G2.

**Figure 4.**
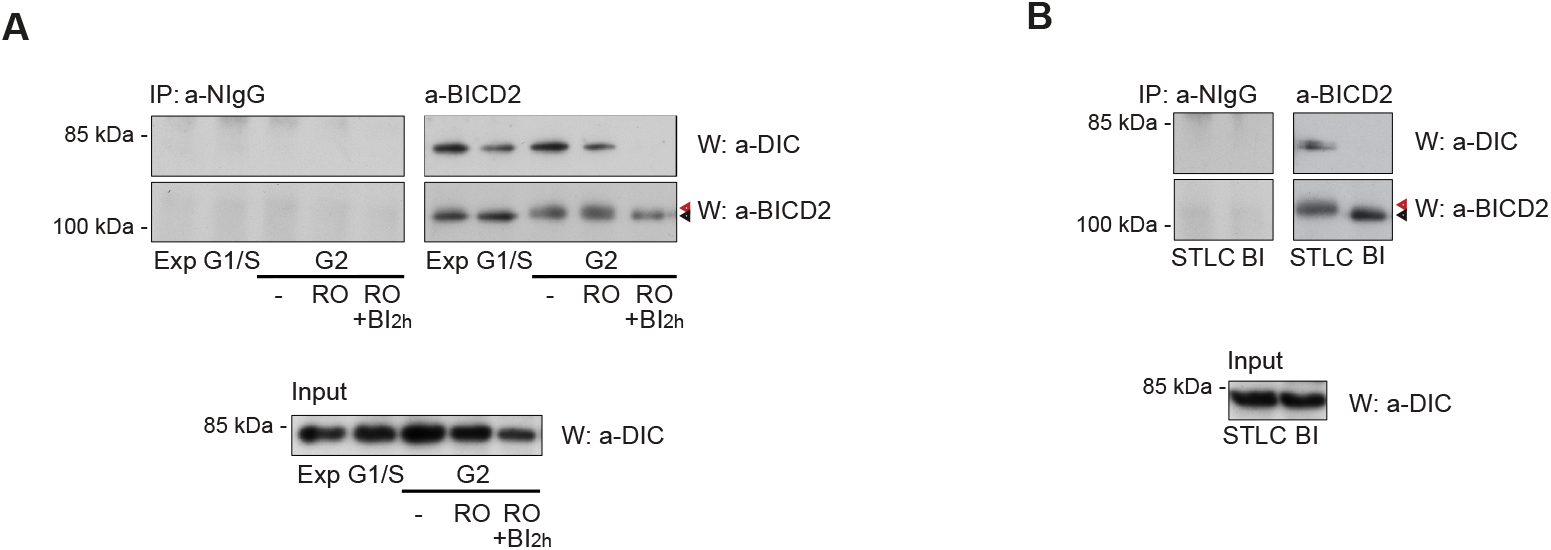
BICD2 forms a complex with dynein in G2/M that is dependent on PLK1 activity. **(A)** PLK1 inhibition disrupts the interaction between BICD2 and dynein in G2. Normal IgG (*NIgG*) or anti-BICD2 immunoprecipitates from HeLa cells at different phases of the cell cycle were analyzed by western blot using anti-DIC or anti-BICD2 antibodies. DIC levels in the corresponding extracts are shown in the lower panel. *Exp*, exponentially growing cells; *G1/S*, cells arrested at the G1/S border after a double thymidine block; *G2*, cells in G2 (*-*, untreated G2 cells, 8h after being released form a double thymidine block; *RO*, RO-3306-arrested G2 cells (9 M, 16h); *RO+BI*, RO-3306-arrested G2 cells treated for 2h with 100 nM BI2536). Cell cycle assignation was confirmed by FACS. Note the increased apparent molecular weight of BICD2 in G2 cells with undisturbed PLK1 activity, arrowheads. **(B)** PLK1 inhibition also disrupts the interaction between BICD2 and dynein in mitosis. As in (A). *STLC*, cells arrested in prometaphase with STLC (5 μM, 16h); *BI*, cells arrested in prometaphase with BI2536 (100 nM, 16h). Note the increased apparent molecular weight of BICD2 in STLC- but not in BI2536-arrested cells, arrowheads.

To further assess the importance of PLK1 activity controlling the formation of BICD2/dynein complexes we chronically inhibited the kinase with BI 2536, a treatment that results in prometaphase arrest (Figure 4B). BICD2 immunoprecipitates from cells treated with the PLK1 inhibitor were not associated with any observable amount of dynein. In contrast, treatment of the cells with STLC, an inhibitor of the kinesin-5 Eg5/KIF11 that equally arrests cells in prometaphase (Skoufias et al., 2006), did not interfere with the ability of dynein to coimmunoprecipitate with BICD2. Of note, in G2 and early mitosis BICD2 shows a high apparent molecular weight that is not observed when the activity of PLK1 is inhibited. (Figure 4A and 4B, arrowheads), suggesting that the kinase is responsible for a significant part of the phosphorylation of BICD2 in these phases of the cell cycle. Finally, acute treatment of exponentially growing cells with BI 2536, RO-3306 or even the pan-CDK inhibitor roscovitine did not interfere significantly with the amount of dynein bound to BICD2 (Supplementary Figure S4). We interpret these results as suggesting that outside of G2 the interaction of BICD2 with the dynein motor complex is independent of CDK/PLK1 phosphorylation, being regulated through the action of other activating mechanisms (e.g. binding to Rab6 or other cargo, or phosphorylation by other protein kinases).

### BICD2 and dynein recruitment to the nuclear envelope in G2 is dependent on PLK1 activity and BICD2 Ser102 modification

We finally aimed to reveal the functional significance of BICD2 phosphorylation in the context of the physiological roles of the adaptor in G2/M. For this we studied how the activity of PLK1 affected the recruitment of BICD2 and dynein to the nuclear envelope. As expected from previously published work (Splinter et al., 2010; Baffet et al., 2015), essentially all control HeLa cells in G2 (with a strong cyclin B1 cytoplasmic staining) showed BICD2 and dynein (DIC) at the nuclear envelope (Figure 5A and B). Most of them had high perinuclear levels of both proteins. Incubation with the pan-CDK inhibitor roscovitine (Meijer et al., 1997) strongly reduced both BICD2 and dynein staining at the nuclear envelope as described before (Baffet et al., 2015). RO-3306, which selectively inhibits CDK1 (>10-fold selectivity vs. CDK2, (Vassilev et al., 2006), only partially displaced BICD2 and dynein from the nuclear envelope, consistent with the notion that CDK2 could have an important role alongside CDK1 in the regulation of the protein complexes in G2. Importantly, PLK1 inhibition with BI 2536 had a strong effect on both BICD2 and dynein localization in G2, resulting in more than 50% of cyclin B-positive cells with no apparent BICD2 or dynein localization at the nuclear envelope. In addition, most cells that retained some BICD2 or dynein perinuclear staining almost entirely had low levels of both the adaptor and the motor at the nuclear envelope. Baffet *et al*. reported similar results for dynein using BTO-1, an alternative PLK1 inhibitor (McInnes et al., 2006; Baffet et al., 2015), and previously Li *et al*. had reported that PLK1 inhibition with BI 2536 interferes with p150 localization to the envelope (Li et al., 2010), but to our knowledge this is the first time that the activity of PLK1 is shown to regulate BICD2 recruitment to the nuclear envelope.

**Figure 5.**
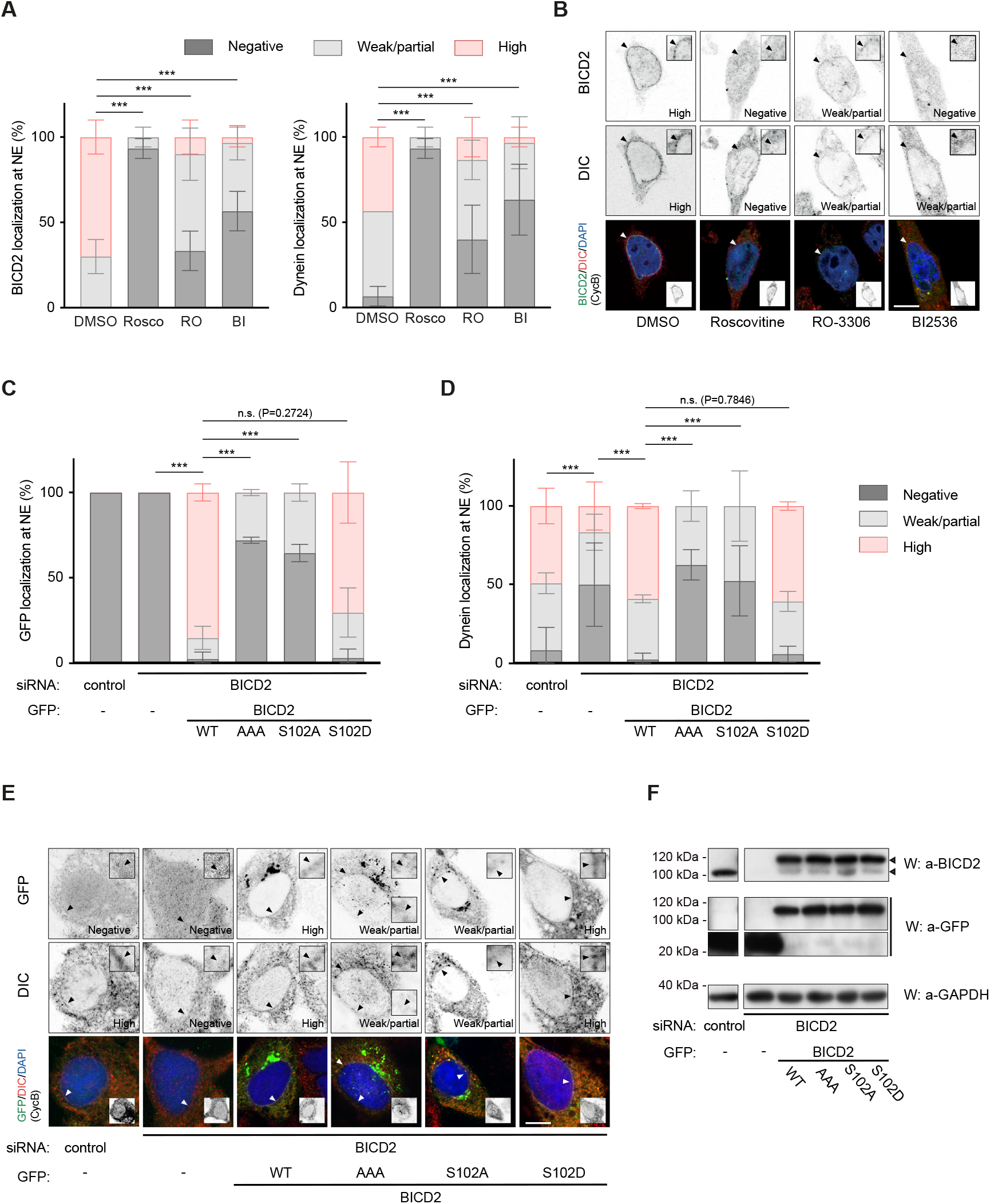
PLK1 activity and BICD2 Ser102 phosphorylation are necessary for normal BICD2 and dynein localization at the nuclear envelope in G2. **(A-B)** CDK and PLK1 inhibition strongly interfere with BICD2 and dynein localization to the nuclear envelope in G2 cells. **(A)** HeLa cells treated with DMSO, 55 μM Roscovitine (*Rosco*), 9 μM RO-3306 (*RO*) or 100 nM BI2536 (*BI*) for 1h were immunostained with the indicated antibodies plus DAPI. For better visualization of NE proteins, nocodazole (10 µM) was included in the last 30 min prior to fixation. G2 cells, identified by positive cyclin B1 cytoplasmic staining, were scored for BICD2 and DIC at the nuclear envelope and the results quantified as shown (n=3 biological replicates, 10 cells per experiment; mean ± SD is shown; statistical significance was analyzed using a Chi square test). **(B)** Example cells for each condition are shown. Scale bar, 10 µm. **(C-F)** Phosphomimetic BICD2 S102D, but not BICD2 AAA or BICD2 S102A, is able to rescue dynein NE localization in G2 cells with low endogenous BICD2 levels. HeLa cells were transfected with the indicated siRNAs and after 48h transfected again with siRNAs plus the indicated GFP-tagged cDNA constructs. After 24h cells were fixed and immunostained with the indicated antibodies plus DAPI. For better visualization of NE proteins, nocodazole (10 µM) was included in the last 30 min prior to fixation. GFP-positive G2 cells, identified by positive cyclin B1 cytoplasmic staining, were scored for GFP **(C)** and DIC **(D)** at the nuclear envelope and the results quantified as shown (n=3 biological replicates, 8-15 cells per experiment; mean ± SD is shown; statistical significance was analyzed using a Chi square test). **(E)** Example cells for each condition (scale bar, 10 µm). **(F)** Expression levels of endogenous BICD2 and GFP-fusion proteins of a representative experiment.

We next sought to determine whether these observations resulted from PLK1 phosphorylation of BIDC2 at Ser102. For this we studied the ability of different recombinant forms of BICD2 to rescue the RNAi-mediated downregulation of endogenous BICD2 in G2 HeLa cells (Figure 5C-F). As described (Splinter et al., 2010), depletion of BICD2 strongly reduced the localization of dynein at the nuclear envelope in cyclin B-positive cells (Figure 5D). This effect was reverted upon expression of GFP-BICD2 wild type, which was able to localize to the nuclear envelope (unlike GFP alone, Figure 5C). GFP-BICD2 AAA (not phosphorylatable by CDK1 and thus unable to bind PLK1) and GFP-BICD2 S102A did not rescue endogenous BICD2 depletion, resulting in cells that either lacked dynein or had low levels of the motor at the nuclear envelope (Figure 5D). Both phosphonull mutants showed a disrupted localization to the nuclear envelope in G2 (Figure 5C), suggesting that besides regulating binding to dynein, PLK1 phosphorylation of BICD2 controls at least partially the interaction of the adaptor with the nuclear envelope. Importantly, the phosphomimetic GFP-BICD2 S102D behaved similarly to the wild type form, both localizing to the nuclear envelope and supporting the recruitment of dynein to this site (Figure 5C and D). In fact, the phosphomimetic mutation of Ser102 was able to rescue dynein localization to the nuclear envelope even in a BICD2 form that cannot be phosphorylated by CDK1 (BICD2 [S102D, T319A, S320A, T321A]; supplementary Figure S5A). This mutant, but not BICD2 AAA can also support dynein binding, suggesting that the exclusive role of _319_TST_320_ phosphorylation is to allow recruitment of PLK1 and the subsequent modification of Ser102 (Supplementary Figure S5).

Altogether our results show that PLK1 phosphorylation of BICD2 Ser102 is a key step for controlling dynein recruitment to the nuclear envelope in G2.

### Centrosome tethering to the nucleus in G2 and separation in early mitosis depend on BICD2 Ser102 modification

Recruitment of dynein at the nuclear envelope in G2 and early mitosis results in tethering of centrosomes to the nucleus, a process that coexists with centrosome separation in early prophase. Separation is tightly regulated and depends on the actomyosin network, microtubules, kinesins and dynein (Agircan et al., 2014). We have shown in the past that PLK1 is necessary for normal centrosome separation and that this is at least partially the result of its role in controlling the phosphorylation of the kinesin Eg5 and its localization to the centrosomes (Bertran et al., 2011). We asked whether Ser102 phosphorylation may also have a role in centrosome separation. We have observed the timing of centrosome separation to be variable among different cell types (Eibes et al., 2018). In G2 HeLa cells showed centrosomes that were adjacent or very close to the nucleus but still unseparated (Figure 6A-C). Downregulation of BICD2 resulted in bigger centrosome to nucleus distances, with few cases in which the centrosomes were as far as 15-20 µm away from the nucleus (mean distance of 4.1 µm vs. 1.6 µm in controls, Figure 6A). This was not accompanied by centrosome separation since in all cases the two centrosomes remained in close proximity (Figure 6B). Expression of GFP-BICD2 completely rescued the BICD2 depletion phenotype, with practically all cells showing centrosomes that were adjacent to the nucleus (mean distance of 0.2 µm). Expression of GFP-BICD2 [T319A, S320A, T321A] or GFP-BICD2 S102A only partially rescued centrosome-nucleus distances (mean distances of 2.0 µm and 2.1 µm, respectively). In contrast, expression of GFP-BICD2 S102D resulted in distances that were similar or even smaller to those observed for wild type BICD2 (mean distance of 0.7 µm). These results show that the modification of BICD2 Ser102 is involved in controlling centrosome tethering to the nucleus in G2.

**Figure 6.**
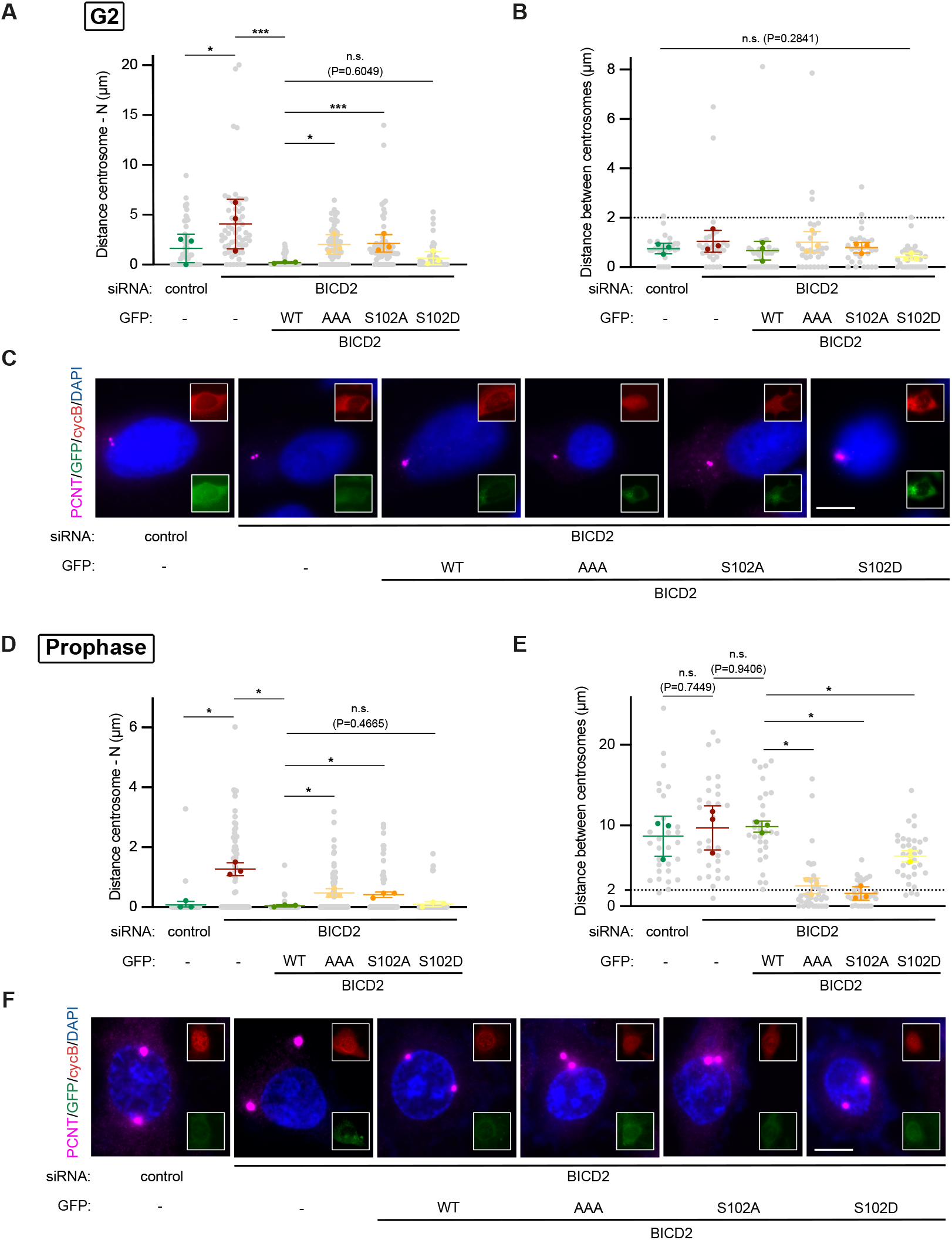
BICD2 Ser102 phosphorylation regulates centrosome tethering to the nucleus and separation during G2 and early mitosis. **(A-C)** Phosphomimetic BICD2 S102D, but not BICD2 AAA or BICD2 S102A, is able to rescue centrosome tethering to the nucleus (*N*) in G2 cells with low endogenous BICD2 levels. HeLa cells were transfected with the indicated siRNAs and cDNAs as in Figures 5C-F, fixed and immunostained with the indicated antibodies plus DAPI. Pericentrin was used as a centrosomal marker. **(A)** Centrosome to nucleus and **(B)** centrosome to centrosome distances were quantified in GFP-positive G2 cells (identified by positive cyclin B1 cytoplasmic staining; n=3 biological replicates, 10 cells per experiment; individual replicate means plus mean of replicates ± SD are shown; statistical significance analyzed using one-way ANOVA with post hoc analysis). **(C)** Example cells for each condition are shown. Scale bar, 10 µm. Centrosomes separated by more than 2µm (dashed line) are considered to be separated. **(D-F)** Phosphomimetic BICD2 S102D, but not BICD2 AAA or BICD2 S102A, is able to rescue centrosome separation in prophase cells with low endogenous BICD2 levels. HeLa cells were transfected, fixed and stained as above. Centrosome to nucleus (*N*) **(D)** and centrosome to centrosome **(E)** distances were quantified in GFP-positive prophase cells (identified by apparent chromosome condensation as assessed with DAPI staining, intact nuclei as assessed by the shape of the DNA and the apparent exclusion of cytoplasmic markers from the nucleus, plus the appearance of nuclear cyclin B1 staining ; n=3 biological replicates, 10 cells per experiment; individual replicate means plus mean of replicates ± SD are shown; statistical significance analyzed using one-way ANOVA with post hoc analysis). **(F)** Example cells for each condition are shown. Scale bar, 10 µm. Centrosomes separated by more than 2µm (dashed line) are considered to be separated.

We next inquired whether the failure to modify Ser102 would also result in abnormal distance between centrosomes at the beginning of mitosis, when centrosomes separate in HeLa cells (Figures 6D-F). For this we studied prophase cells, identified by their condensed chromosomes and intact nuclei, and presence of nuclear cyclin B1 (independent experiments showed that these features paralleled histone H3[Ser10] phosphorylation in all cases, confirming correct cell cycle phase staging). Almost all control prophase cells had separated centrosomes that were adjacent to the nucleus (8.8 µm mean distance between centrosomes; we considered centrosomes to be separated when the intercentrosomal distance was more than 2µm). Downregulation of BICD2 detached centrosomes from the nucleus as in G2 (Figure 6D). The organelles were separated (mean intercentrosomal distance of 9.8 µm, Figure 6E) with a distribution in the cytoplasm that appeared to be random. GFP-BICD2 wild type was able to completely rescue centrosome attachment to the nucleus, with the organelles tethered to the nuclear membrane and separated (mean distance of 10.0 µm). Both GFP-BICD2 AAA or GFP-BICD2 S102A were only able to partially rescue centrosome attachment to the nucleus. Strikingly, both mutants almost completely failed to support centrosome separation showing mean intercentrosomal distances of 2.6 µm and 1.7 µm, respectively. We conclude that BICD2 phosphorylation by PLK1 is necessary for the recruitment of normal amounts of (active) dynein to the nuclear envelope to promote centrosome separation. Indeed, GFP-BICD2 S102D, which was observed to completely rescue centrosome attachment to the nucleus in both G2 and prophase (Figures 6A and D), allowed for close to normal centrosome separation in this early phase of mitosis (mean intercentrosomal distance of 6.3 µm, Figures 6B and E).

Altogether our results suggest that the modification of Ser102 is necessary for the recruitment of BICD2 and dynein to the nuclear envelope and for centrosome tethering to the nucleus in G2 and early M, as well as for normal centrosome separation in prophase. They highlight the importance of PLK1 phosphorylation of the dynein adaptor in these phases of the cell cycle and, moreover, identify adaptor phosphorylation as a previously unknown regulatory mechanism for recruitment and activation of the dynein machinery.

## Discussion

Dynein multiple functions suggest that the motor is subject to complex regulation. This likely involves different molecular mechanisms acting in response to a variety of signaling inputs. For instance, dynein itself may be directly regulated through post-translational modification of its different chains (see below). Moreover, associated proteins such as dynactin, LIS1 and NDE1/NDEL1, and a number of structurally diverse cargo adaptors play a major role in modulating the activity of the motor complex (Reck-Peterson et al., 2018; Canty et al., 2021). A number of adaptors are particularly important in this regard. Apart from connecting the dynein complex to specific cargos and other motors such as kinesins, they share the ability to facilitate the interaction between dynein and dynactin, effectively functioning as dynein activators (Reck-Peterson et al., 2018; Olenick and Holzbaur, 2019). Despite their importance, how the different adaptors and their dynein-activating function are regulated when cellular physiology demands has remained obscure. One possibility is that dynein activation is triggered through cargo binding; in this manner motor activity would be spatiotemporally linked to cargo availability. Indeed this regulatory mechanism has been suggested for BICD family members (Hoogenraad and Akhmanova, 2016). Both BICD1 and BICD2 are autoinhibited through the intramolecular interaction between their N- and C-terminal regions that is thought to physically impede the binding of dynein and dynactin (Hoogenraad et al., 2001; Splinter et al., 2012; Terawaki et al., 2015). Binding of the small GTPase Rab6 to the C-terminus of BICD1/2 liberates the N-terminus, allowing it to organize an active motor complex with dynein and dynactin in order to transport exocytotic lipid vesicles (Hoogenraad et al., 2001; Matanis et al., 2002; Huynh and Vale, 2017). This does not seem to be the case for other adaptors such as BICDL1 (Urnavicius et al., 2018) or HOOK proteins (Olenick et al., 2016) which apparently interact with dynein and dynactin independently of cargo binding.

In this work we put forward adaptor “activation” by phosphorylation as an alternative mechanism of assembling active dynein complexes with the potential to increase the complexity and flexibility of dynein regulation. Indeed, our results show that in response to phosphorylation by the G2/M protein kinase PLK1 BICD2 is able to bind dynein as well as dynactin and form an active motor complex. In fact, dynein binding to BICD2 is almost completely dependent on the activity of PLK1 in G2 and M (but not other phases of the cell cycle), thus indicating that phosphorylation is the prevalent mechanism regulating the adaptor in these phases of the cell cycle. Considering the observed changes in BICD2 electrophoretic mobility as a proxy for phosphorylation, BICD2 seems to be phosphorylated exclusively in G2/M, when PLK1 is active (Bruinsma et al., 2012; Zitouni et al., 2014). In the light of previous findings, our data suggests that the adaptor could be subject to dual regulation. During in G1 and S, in the context of lipid vesicle transport, BICD2 may be controlled by the previously described Rab6-based mechanism. And in G2 and M, when a major reorganization of the intracellular membrane system occurs and BICD2 switches partner binding to RanBP2 at the nuclear pores, by PLK1.

Our results suggest that the regulation of BICD2 during this switch does not depend exclusively on PLK1, but also on 2/M cyclin dependent kinases such as CDK1. CDKs regulate PLK1 binding to BICD2 by modifying a specific motif in BICD2 (_319_TST_321_ in mice, _317_TST_319_ in humans) that is recognized by PLK1 PBD, a specialized phosphoserine/phosphothreonine binding domain involved in targeting the kinase to its substrates (Elia et al., 2003b). We therefore propose that two kinases regulate BICD2 in a concerted manner, with CDK1 (or possibly CDK2, see below) acting as a “priming” kinase that allows BICD2 to interact with PLK1, ultimately resulting in the phosphorylation of the adaptor by PLK1 and its “activation”. Although *in vitro* and possibly in conditions of excess PLK1 activity this priming step might not be required (e.g., we observed *in vitro* phosphorylation of BICD2 fragments by PLK1 independently of a priming phosphorylation), our results suggest that in cells it is a necessary step for BICD2 regulation by PLK1, since mutation of the CDK sites (BICD2 [T319A, S320A, T321A]) is similarly deleterious to G2/M functions of BICD2 as mutation of the main PLK1 site (BICD2 S102A). Regarding the role of CDK1, we observed that cells arrested with RO-3306, a specific inhibitor of the kinase, still show sizeable (although slightly reduced) levels of BICD2/dynein complexes. This suggests that either other CDK family members active in G2 (i.e. CDK2) could have a role as PLK1 priming kinases. Alternatively, a long arrest with high levels of PLK1 may result in BICD2 phosphorylation independently of its binding to CDK1 (see for example (Loncarek et al., 2010)). Further experiments will be needed to clarify this point.

PLK1 rapidly phosphorylates BICD2 at Ser102, a modification that our results show has major structural and functional consequences. Ser102 is located at the predicted elongated N-terminal coiled-coil region, close to the dynein-interacting CC1 box (see Figure 1H). Notably, in humans, a mutation of the nearby Ser107, found in a rare form of spinal muscular atrophy (SMALED2; OMIM 615290) results in an enhanced ability to form motile dynein complexes (Huynh and Vale, 2017).

Electron micrographs of wild type BICD2 showed that the molecule is predominantly in a triangular or looped conformation, similar to what has been described for *Drosophila* BICD (Stuurman et al., 1999; Sladewski et al., 2018), although in the mammalian homolog the looped part seems to comprise a bigger part of the molecule. We propose that this triangular shape corresponds to the hypothesized “closed” conformation of the adaptor, resulting from the intramolecular interaction between its N- and C-terminal parts. (Hoogenraad et al., 2001). Importantly, the modification of Ser102 seems to favor a conformational change in BICD2, resulting in BICD S102D being mostly in an extended rod-shaped “open” conformation, possibly more amenable to interact with dynein/dynactin. Accordingly, DeepMind-based methods for protein structure prediction (Jumper et al., 2021) open the possibility that BICD2 Ser102 could be at the interaction interface between the CC1 and the CC3 coiled-coil regions. We cannot discard that the differences found in the micrographs of WT and S102D BICD2 could be caused by changes in the way the molecules interact with the support film, the mutation favoring side views of the molecules. Although we think this is less likely, even in this case, these effects would be a consequence of modifications in the structure of the protein. In summary, EM shows that the phosphomimetic S102D mutation induces conformational changes in BICD2, which we propose could result in a more extended or “open” BICD2 conformation. Future efforts aimed at determining the atomic structure of both wild type and S102D BICD2 should reveal the molecular basis of this conformational change, be it electrostatic or steric repulsion or a registry shift (Noell et al., 2019). But altogether our interpretation that the phosphorylation of Ser102 results in the functional opening of BICD2, is also supported by biochemical data and cell-based assays. We show that it disrupts the autoinhibitory intramolecular interaction between the C- and the N-terminal regions of the dimeric structure of the adaptor, favoring BICD2 binding to dynein and dynactin and the formation of active motor complexes that accumulate at the minus ends of microtubules.

Since we propose that BICD2 phosphorylation is the main regulatory mechanism for this adaptor in G2/M, we have studied its relevance during the recruitment of dynein to the nuclear envelope in these phases of the cell cycle by using both protein kinase inhibitors and different BICD2 mutants. Dynein nuclear envelope recruitment is important for the intracellular positioning of nucleus and centrosomes as well as for centrosome separation in late G2 and early M (Gönczy et al., 1999; Robinson et al., 1999; Splinter et al., 2010; Bolhy et al., 2011; Raaijmakers et al., 2012). Additionally it has a key role driving the specialized apical nuclear migration in neural progenitors (Hu et al., 2013). This process, although not studied here, may also involve regulation through phosphorylation of BICD2, since it occurs at the same cell cycle stage and also involves nuclear envelope-localized dynein. We observed that in HeLa cells dynein recruitment to the nuclear envelope in G2 depends on both the activity of CDK1 and PLK1. CDK1 inhibition is not as disruptive as pan-CDK (or PLK1) inhibition thus suggesting the additional involvement of a second G2 CDK (i.e. CDK2). CDK1 has been previously reported to control dynein recruitment to the nuclear envelope though its ability to phosphorylate RanBP2, CENP-F and NDE1 and thus regulate the two known pathways of dynein nuclear recruitment (Baffet et al., 2015; Wynne and Vallee, 2018). Our results add an additional independent layer of regulation that acts directly on the BICD2 adaptor. They are not only in agreement with the previously proposed involvement of PLK1 (Baffet et al., 2015), but place the Polo-family kinase in a central position during the regulation of dynein nuclear envelope recruitment.

Remarkably, phosphorylation of BICD2 by CDKs at _319_TST_321_ and by PLK1 at Ser102 does not only control the ability of the adaptor to bind dynein and recruit it to the nucleus, but also the nuclear recruitment of BICD2 itself. This strongly suggests that BICD2 needs to be phosphorylated and presumably in the “open” conformation to be able to bind to its nuclear receptor, RanBP2. In this view RanBP2 would not behave as a “canonical” cargo such as Rab6, able to open the adaptor upon binding and allowing it to interact with dynein. Instead, it would be the phosphorylation of BICD2 Ser102 that would change BICD2 conformation, allowing the adaptor to bind both dynein *and* RanBP2 (note that directly binding of unmodified BICD2 to RanBP2 has only been shown *in vitro* for the C-terminal region of the adaptor (Splinter et al., 2010; Baffet et al., 2015). The fact that RanBP2 and Rab6 have different binding sites in BICD2 (Schlager et al., 2010; Terawaki et al., 2015), could possibly provide a molecular explanation for this hypothesis, which surely needs to be further explored. The necessity of a regulated interaction between BICD2 and RanBP2 in G2 may be understood in view of the reported affinity between the C-terminus of BICD2 and RanBP2, which is much higher than the affinity for Rab6(GTP) (Noell et al., 2018). One can speculate that unregulated high-affinity binding of RanBP2 to BICD2 would result in massive recruitment of the adaptor to the nuclear pore. In fact Noell *et al*. calculated that ∼1000 BICD2 molecules per cell would be recruited to the nucleus by RanBP2 in the absence of regulation (Noell et al., 2018). Therefore CDK1 and PLK1 phosphorylation and opening of BICD2 ensures that the adaptor interacts with the nucleoporin exclusively at the right cell cycle phase, something that will be further favored by CDK1 phosphorylation of RanBP2 (Baffet et al., 2015).

Using different mutants, we find that BICD2 phosphorylation controls centrosome tethering to the nucleus and centrosome separation in G2 and early mitosis. We show that BICD2 downregulation significantly increases centrosome to nucleus distances in both G2 and prophase, and that expression of wild type BICD2 is able to rescue this phenotype. This is similar to what has been reported for prophase U2OS cells (Splinter et al., 2010), but in contrast to that observed in prophase HeLa cells (Raaijmakers et al., 2012). The differences may be due to variations in the time of activation or the strength of the later acting Nup133/CENPF/NDE1/NDEL1 dynein recruiting pathway (Bolhy et al., 2011) in the specific cell lines used in each study. For example, we observed centrosomes to be more closely tethered to the nucleus in prophase than in G2, which may be attributed to the additional activity of this later acting pathway. Both in G2 and prophase the disruption of BICD2 phosphorylation by CDK1 and PLK1 interfere with the ability of the adaptor to support normal centrosome tethering to the nucleus. In contrast, the phosphomimetic BICD2 S102D rescues centrosome to nucleus distances almost as well as the wild type form, confirming that phosphorylation of this residue ultimately regulates this process.

We similarly implicate BICD2 and its phosphorylation in the control of centrosome separation. Interestingly, while dynein participates in centrosome separation in both *Drosophila* and *C.elegans* (Gönczy et al., 1999; Robinson et al., 1999), the motor has been reported to be dispensable for the separation of the organelles in prophase in U2OS cells with normal kinesin Eg5 levels (Tanenbaum et al., 2008; Splinter et al., 2010; Raaijmakers et al., 2012). In RPE-1 and U2OS cells BICD2 also seems not to be necessary (Raaijmakers et al., 2012). According to this, the role of dynein during centrosome separation seems to be to keep the centrosomes close to the nucleus, antagonizing (possibly together with perinuclear actin (Stiff et al., 2020) the forces produced by Eg5, that push centrosomes far apart but also away from the nucleus. We have indeed observed that the downregulation of BICD2 in prophase (but not in G2 where presumably Eg5 is still not active) results in centrosomes that are mostly separated. However, we note that the centrosomes appeared to be randomly distributed in the cytoplasm, similar to what was reported for treatments that disrupt the microtubule cytoskeleton (Jean et al., 1999; Meraldi and Nigg, 2001). This may suggest that centrosomes were not be actively separating but passively drifting apart. Importantly, we find that BICD2 seems to be having a previously unreported role during separation, as the expression of mutant forms of the adaptor that cannot be phosphorylated by CDK or PLK1 results in small intercentrosomal distances. In contrast expression of BICD2 S102D not only rescues centrosome tethering to the nucleus but also centrosome separation to an extent that is also observed for the wild type form of the adaptor. The elucidation of the molecular basis of these observations is beyond the scope of this project, although our preliminary results show that BICD2 is involved in the recruitment of Eg5 to the centrosomes during prophase, an observation that may be the basis of the observed phenotype. Nonetheless, our results add a new regulatory node to the convolutedly controlled process of (S.Eibes and J. Roig, unpublished data), a process that is important for the normal segregation of chromosomes later in mitosis (Silkworth et al., 2012). We and others have previously shown that centrosome separation depends on Eg5 phosphorylation by CDK1 and NEK6/7, the latter kinases activated by the related NEK9 downstream of PLK1 and also CDK1 (Blangy et al., 1995; Bertran et al., 2011). We can now add CDK1 and PLK1 phosphorylation of BICD2 to this regulatory network.

Our work shows that the formation of active dynein complexes can be regulated not only through cargo binding to specific dynein adaptors, but also through adaptor phosphorylation. Further studies should establish whether other adaptors besides BICD2 share this regulatory mechanism. This may be the case for adaptors relying on autoinhibition for the control of their activity, such as the closely related BICD1 or *Drosophila* BICD. In the case of BICD1 the GSK-3ß kinase is in fact able to modify its C-terminus in the context of the regulation of centrosomal components and aster microtubule dynamics in interphase (Fumoto et al., 2006). And BICD is known to depend on phosphorylation for its roles during oocyte differentiation, although the details of this are unknown (Suter and Steward, 1991). Our results establish that phosphorylation has a central role in regulating dynein physiology. Indeed, not only cargo adaptors but most dynein subunits are phosphorylated *in vivo* (e.g. see (Dillman and Pfister, 1994), and https://www.phosphosite.org/), possibly by a number of protein kinases including PLK1 (Li et al., 2010; Bader et al., 2011). The detailed study of this will surely expand our understanding of the functional complexity of the dynein motor complex.

## Materials and Methods

### Mammalian expression constructs and siRNAs

pEGFP-C2-BICD2, containing the cDNA for mouse BICD2 was a gift from Anna Akhmanova (Utrecht University) and is described in (Hoogenraad et al., 2001). Fragments and mutant forms of BICD2 were produced using QuickChange Lighting Site-Directed Mutagenesis kit (Agilent Technologies) according to the manufacturer’s instructions using the following primers and the appropriate reverse complements:

BICD2 [1-575]: 5’-GAGGGCCGCGGGTGACGGTCACCTGTC-3’;
BICD2 [576-820]: 5’-GATCTCGAGAGAATTCGCCACCCGCCGGTCACCTGTCCTCTTG-3’;
BICD2 [T319A, S320A, T321A]: 5’-CATCCTTGGACAACAAGGCAGCCGCACCCAGGAAGGATGG-3’;
BICD2 S102A: 5’-GAGAGCCGGGAGGAGGCCCTGATCCAGGAGTCG-3’;
BICD2 S102D: 5’-GAGAGCCGGGAGGAGGACCTGATCCAGGAGTCG-3’.

To determine the mitochondrial clustering ability of the different BICD2 forms, the desired BICD2 cDNAs were amplified from pEGFP constructs by PCR using Phusion High-Fidelity DNA polymerase (Thermo Fisher) and inserted using a Gibson assembly strategy (NEB) into a pEGFP-C1-MTD plasmid, also containing the cDNA of the isoform G of centrosomin from *Drosophila melanogaster* (obtained from the Drosophila Genomics Resource Center, accession number AT09084). pEBG-BICD2 [272-540] and pEBG-BICD2 [541-820] were obtained by subcloning the corresponding cDNAs, amplified by PCR using Pfu Ultra HF DNA polymerase (Agilent Technologies) and pEGFP-C2-BICD2 as a template, into empty pEBG vector. Coding regions of all constructs was sequenced prior to use.

For endogenous BICD2 knockdown we used the following siRNA sequence, after (Splinter et al., 2010): 5’-GGAGCUGUCACACUACAUGUU-3’. A siRNA against luciferase with the following sequence was used as control: 5’-CGUACGCGGAAUACUUCGA-3’.

### Bacterial expression

pGEX-PLK1 PBD [345–603] was previously described (Bertran et al., 2011). pGEX-BICD2 [1-271] and pGEX-BICD2 [272-540] were obtained by subcloning the corresponding cDNAs, amplified by PCR using Pfu Ultra HF DNA polymerase and pEGFP-C2-BICD2 as a template, into empty pGEX-4T1. pGEX-BICD2 [272-540; T319A, S320A, T321A] mutant was produced using QuickChange Lighting Site-Directed Mutagenesis kit according to the manufacturer’s instructions with the primers described above. Proteins were expressed in *E. coli* BL21(DE3) pLysS cells and purified using glutathione agarose beads (GE Healthcare) according to the manufacturer’s instructions. BICD2 fragments were eluted with 10 mM reduced L-glutathione (Merck Life Science). PLK1 PBD was kept bound to the agarose and stored at 4°C for subsequent pull-down experiments.

### Baculovirus expression

pLIB-TwinStrep-3C-BICD2 was constructed using a Gibson assembly strategy with the corresponding BICD2 cDNA (amplified from pEGFP constructs by PCR using Pfu Ultra HF DNA polymerase), plus pOPINN1S3CTwinStrep-3C and pLIB plasmids. pLIB-TwinStrep-3C-BICD2 S102D was made using QuickChange Lighting Site-Directed Mutagenesis kit.

Full length BICD2 proteins were produced by baculovirus-mediated expression in insect cells. Bacmids were generated by Tn7 transposition of pLIB-TwinStrep-3C-BICD2 constructs into the EMBacY baculovirus genome as described (Weissmann and Peters, 2018). Baculoviruses were generated as described (Scholz and Suppmann, 2017) with minor modifications.

Full length BICD2 WT and BICD2 S102D were purified from infected cells using the N-terminal TwinStrep-tag. Frozen pellets were resuspended on ice in Lysis Buffer B (50 mM Tris pH 8.0, 300 mM NaCl, 2 mM β-mercaptoethanol, 0.1% IGEPAL CA-630, 2x complete protease inhibitor cocktail EDTA-free (Roche), 5% glycerol) and lysed in a Dounce tissue grinder with 20 strokes on ice. After lysis, crude extracts were supplemented with 10 µl DNArase (c-Lecta), incubated 5 min on a tube roller mixer at 4°C, and centrifuged for 25 min at 20,000g at 4°C. Cleared extracts were supplemented with 2 mg of avidin (E-proteins) and bound by gravity flow to 1 ml of StrepTactinXT 4Flow high capacity (IBA) equilibrated in Wash Buffer B (25 mM Tris pH 8.0, 300 mM NaCl, 2 mM β-mercaptoethanol + 5% glycerol). Resin was washed with 10 column volumes of Wash Buffer B and eluted with Elution Buffer B (Wash Buffer B + 50 mM biotin). Peak fractions containing BICD2 were identified by SDS-PAGE, pooled, concentrated to 300-350 µl using Vivaspin 6 30,000 MWCO concentrators (Sartorius), and loaded onto a Superdex 200 10/300 GL column equilibrated in SEC Buffer (25 mM Tris pH 8.0, 300 mM NaCl, 0.5 mM TCEP+ 5% glycerol). Peak fractions were pooled, concentrated with Vivaspin 500 30,000 MWCO concentrators to 0.9-2 mg/ml, aliquoted, snap-frozen in liquid nitrogen and stored at - 80°C until further use.

### Protein phosphorylation and phosphosite determination

*In vitro* protein kinase assay was carried out by incubation of 1 ug purified protein substrates in the presence or absence of 100 ng PLK1 (Thermo Fisher) or CDK1/cyclin B1 (Thermo Fisher) in Phosphorylation Buffer (50 mM MOPS pH 7.4, 5 mM MgCl_2_, 10 mM β-glycerophosphate, 1 mM EGTA, 1mM DTT plus 100 µM ATP) at 25 °C for the indicated times. Phosphate incorporation was visualized by adding 1 µCi ^32^P-γ-ATP (200 cpm/pmol, Perkin Elmer), plus autoradiography or a PhosphorImager system (Molecular Dynamics). Quantifications were done using the PhosphorImager system. Non-radioactive reaction duplicates were used for phosphosite determination. After SDS-PAGE, relevant bands were cut, digested with trypsin or, when indicated, chymotrypsin and analyzed by nano-LC-MS/MS using an Orbitrap Fusion Lumos TM Tribrid instrument at the IRB Barcelona Mass Spectrometry and Proteomics Core Facility. Samples were run through a PepMap100 C18 u-precolumn (Thermo Fisher) and then separated using a C18 analytical column (Acclaim PepMap RSLC, Thermo Fisher) with a 90 minutes run, comprising three consecutive steps with linear gradients from 3 to 35% B in 60 minutes, from 35 to 50% B in 5 minutes, and from 50 % to 85 % B in 1 minute, followed by isocratic elution at 85 % B in 5 minutes and stabilization to initial conditions (A= 0.1% FA in water, B= 0.1% FA in CH_3_CN). The column outlet was directly connected to an Advion TriVersa NanoMate (Advion) fitted on an Orbitrap Fusion Lumos™ Tribrid (Thermo Scientific). The mass spectrometer was operated in a data-dependent acquisition (DDA) mode. Survey MS scans were acquired in the orbitrap with the resolution (defined at 200 m/z) set to 120,000. The lock mass was user-defined at 445.12 m/z in each Orbitrap scan. The top speed (most intense) ions per scan were fragmented in the HCD cell and detected in the orbitrap. The ion count target value was 400,000 and 50,000 for the survey scan and for the MS/MS scan respectively. Target ions already selected for MS/MS were dynamically excluded for 30s. Spray voltage in the NanoMate source was set to 1.60 kV. RF Lens were tuned to 30%. Minimal signal required to trigger MS to MS/MS switch was set to 25,000. The spectrometer was working in positive polarity mode and singly charge state precursors were rejected for fragmentation. A database search was performed with Proteome Discoverer software v2.1 (Thermo Scientific) using Sequest HT, Amanda search engines and SwissProt database Mouse release 2016_11; contaminants database and user proteins manually introduced. Searches were run against targeted and decoy databases to determine the false discovery rate (FDR). Search parameters included trypsin enzyme specificity, allowing for two missed cleavage sites, Methionine oxidation, Phosphorylation in serine/threonine/tyrosine and acetylation in N-terminal as dynamic modifications and Carbamidomethyl in cysteine as static modification. Peptide mass tolerance was 10 ppm and the MS/MS tolerance was 0.02 Da. Peptides with a q-value lower than 0.1 and an FDR < 1% were considered as positive identifications with a high confidence level. The PhosphoRS node was used to provide a confidence measure for the localization of phosphorylation in the peptide sequences identified with this modification.

### Cell culture, synchronization and transfection

HeLa and U2OS cells were grown in DMEM media (Thermo Fisher) supplemented with 10% foetal bovine serum (Thermo Fisher), 2mM L-glutamine (Thermo Fisher) and 100 U/mL penicillin/ 100 ug/mL streptomycin (Thermo Fisher).

Mitotic HeLa cells were obtained by mitotic shake off of cells arrested using either 200 ng/mL nocodazole (Merck Life Science), 100 nM BI2563 (Adooq Bioscience) or 5 µM STLC (Merck Life Science) for 16 h. Cells arrested in G2 were obtained after a 16 hours treatment with 9 µM RO-3306 (Enzo Life Sciences). Cells at the G1/S border and progressing through G2 were obtained using a double thymidine double block. Briefly, cells were plated with media containing 2mM thymidine (Merck Life Science) during 16 hours, washed and released in fresh media. After 8 hours post-release media was supplemented again with 2 mM thymidine during 16 hours. Cells collected at this point were in G1/S. To obtain G2 cells, after the double thymidine block cells were washed, released in fresh media and collected after 8 hours. Cell cycle phase was confirmed by FACS in all cases.

For immunoprecipitation experiments, cells were transfected using linear polyethylenimine (Polysciences Inc.) according to manufacturer’s instructions. For immunofluorescence, either Lipofectamine 2000 (plasmids) or Lipofectamine RNAiMAX (siRNAs) was used, according to manufacturer’s instructions (Thermo Fisher)

### Antibodies

Primary antibodies used were as follows:

a-PLK1 (mouse monoclonal IgG2b, Calbiochem #DR1037)
a-BICD2 (rabbit polyclonal, Abcam #ab117818 for western blot (WB) and immunoprecipitations (IP); for immunofluorescence (IF), rabbit polyclonal #2293, a gift from Anna Akhmanova (Utrecht University), described in (Hoogenraad et al., 2001).)
a-GFP (mouse monoclonal IgG2a, Thermo Fisher #A11120; rabbit polyclonal Torrey Pines #TP401 (IF) and Santa Cruz #sc-8334 (IF))
a-p150 (mouse monoclonal IgG2b, Santa Cruz #sc-365274)
a-DIC (mouse monoclonal IgG2b, Santa Cruz #sc-13524 forw WB and IP; mouse monoclonal IgG2b, Thermo Fisher #14- 97772-80 for IF)
a-GST (mouse monoclonal IgG1, Sigma #SAB4200237)
a-GAPDH (mouse monoclonal IgG1, Santa Cruz #sc-47724)
a-Tom20 (mouse monoclonal IgG2a, Santa Cruz #sc-17764)
a-PCNT (rabbit polyclonal, Abcam #ab4448)
a-CycB (mouse monoclonal IgG1, Santa Cruz #sc-245)

Secondary antibodies used were as follows (Alexa Fluor conjugates, Thermo Fisher; HRP conjugates, R&D):

Alexa Fluor 647 goat anti-rabbit IgG (H+L) #A21244
Alexa Fluor 568 goat anti-mouse IgG2b #A21144
Alexa Fluor 488 goat anti-mouse IgG1 #A21121
Alexa Fluor 488 goat anti-rabbit IgG (H+L) #A11008
Alexa Fluor 647 goat anti-mouse IgG1 #A21240
Alexa Fluor 488 goat anti-mouse IgG2a #A21131
Alexa Fluor 568 goat anti-mouse IgG1 #A21124
Alexa Fluor 647 goat anti-rabbit IgG (H+L) #A21244
Goat a-mouse IgG-HRP #HAF007
Goat a-rabbit IgG-HRP #HAF008

### Immunoprecipitation, pulldowns and western blot

Cells were lysed in Lysis Buffer B2 (20 mM Tris-HCl pH 8, 100 mM KCl, 1% Triton X-100, 10 mM β-glycerophosphate, 25 nM calyculin A, 0.5 mM PMSF, 1 ug/mL aprotinin and 1 ug/mL leupeptin), adapted from (Splinter et al., 2012). To study coimmunoprecipitation of BICD2 with dynein, we used Lysis Buffer D (25 mM HEPES pH 7.4, 50 mM potassium acetate, 2 mM magnesium acetate, 0.2% Triton X-100, 1 mM EGTA, 50 mM NaF, 10 mM β-glycerophosphate, 10 % glycerol, 0.5 mM Mg-ATP, 1 mM DTT, 25 nM calyculin A, 0.5 mM PMSF, 1 ug/mL aprotinin and 1 ug/mL leupeptin), adapted from (Redwine et al., 2017)). Immunoprecipitations and western blotting were performed using standard protocols as described in (Roig et al., 2002) using the respective lysis buffers as wash buffer. For immunoprecipitation protein G coupled to Dynabeads (Thermo Fisher) was used. GST pulldowns were done as described in (Bertran et al., 2011) using glutathione agarose beads (GE Healthcare).

### Immunofluorescence

Cells were grown on Poly-L-Lysine coated coverslips to sub-confluence. After the indicated treatments, cells were rinsed in cold PBS and fixed with methanol at −20 °C for at least 15 minutes. After three washes in PBS, fixed cells were incubated for 20 minutes in Blocking Buffer (3% BSA, 0.1% Triton X-100, 0.02% NaN_3_ in PBS) and stained using the indicated primary antibodies diluted in blocking solution for 1 hour at room temperature. Secondary antibodies were diluted in blocking solution together with DAPI (0.01 mg/mL) (Merck Life Science) and incubated for 20 minutes at room temperature in a dark, humid chamber. Slides were mounted in ProLong Gold Antifade Reagent mounting media (Thermo Fisher), sealed and stored in a cold dark chamber. When indicated, nocodazole (10 µM) was included in the last 30 min prior to fixation to better visualize nuclear envelope proteins (Baffet et al., 2015).

Three channel Z-stack images were acquired in a Leica AF6000 system with an Orca AG camera (Hamamatsu) coupled to a Leica DMI6000B microscope equipped with 63x/1.40NA and 100x/1.40NA oil immersion lens and the standard LAS-AF software followed with deconvolution. Four channel Z-stack images were acquired with a Leica Thunder system with a DMI8 microscope equipped with 100x NA 1.40 HCX PL-APO oil immersion lens and the standard LAS-AF software followed with deconvolution. Four channel Z-stack confocal images were obtained using a Zeiss Lsm780 confocal system with an inverted XYZ Zeiss Axio Observer Z1 microscope equipped with an 63x/1.40NA oil immersion lens and the standard Zeiss software (ZEN).

### Negative staining electronic microscope (EM)

Wild type and S102D BICD2 samples were diluted in Protein Buffer (25 mM Tris-HCl pH 9.0, 300 mM NaCl, 0.5 mM TCEP) to a final concentration of 0.02 mg/ml. Diluted samples were applied onto home-made glow-discharged carbon-coated copper grids and negatively stained with 2% (w/v) uranyl formate, pH 7.0. Electron microscope images were acquired in a FEI Tecnai G2 Spirit microscope with a Lab6 filament and operated at 120 kV, using a TVIPS CCD camera. Microscope images were processed using the cryoSPARC v3.2 (Punjani et al., 2017) software package. Automatically selected BICD2 particles (150,473 BICD2 wild type particles and 106,650 BICD2 S102D particles) were aligned and classified using the reference-free 2D classification tool in cryoSPARC to obtain a clean dataset (30,476 BICD2 wild type particles and 22,021 BICD2 S102D particles).

### Quantification and statistical analysis

Assessment of mitochondria clustering and protein localization at the nuclear envelope, plus measurement of centrosome to nucleus and centrosome to centrosome distances were done using FIJI software. Graphics and statistical analysis were done using Prism 9 software. Details of the number of measurements and biological replicates, as well of the statistical tests used in each case are given in the corresponding figure legends (*p<0.05; ** p<0.01; ***p<0.001).

## Acknowledgements

We are grateful to Anna Akhmanova (Utrecht University) for reagents and comments and Susana Eibes (Danish Cancer Society Research Center) for help with different DNA constructs and comments. We would also like to thank Elena Rebollo and the Molecular imaging Platform at the IBMB-CSIC as well as the Mass Spectrometry & Proteomics Core Facility at the IRB Barcelona for their help and input.

We are grateful to Carme Caelles (UB), Jose Reina (IRB Barcelona) and Martí Aldea (IBMB-CSIC) for helpful comments as members of N.G.-S.’s thesis advisory committee, and Thomas Surrey (CRG), Guillermo de Cárcer (IIB-CSIC) and Neus Agell (UB) for helpful comments as members of N.G.-S.’s PhD committee.

J.R. and N.G.-S.work was funded by the Ministerio de Ciencia e Innovación (MICINN) and the Agencia Estatal de Investigación (AEI) from Spain (MCIN/ AEI/10.13039/501100011033/ FEDER “*Una manera de hacer Europa*”) through Plan Nacional de I+D grants BFU2014-58422-P and PGC2018-096307-B-I00 (to J.R.) and by network grant 2017 SGR 1089 (AGAUR, Generalitat de Catalunya). N.G.-S. was a recipient of FPI Fellowship BES-2015-072446 from MICINN.

Work in the laboratory of J.L. was supported by grants BFU2015-69275-P (MINECO/FEDER), PGC2018-099562-B-I00 (MICINN), network grants 2017 SGR 1089 (AGAUR) and RED2018-102723-T (MICINN), and by intramural funds of IRB Barcelona, supported by CERCA (Generalitat de Catalunya). F.Z. was supported by a fellowship from the “la Caixa” Foundation (ID 100010434, fellowship code LCF/BQ/DI17/11620020) and the European Union’s Horizon 2020 research and innovation program under the Marie Skłodowska-Curie grant agreement no. 713673. J.P. was supported by a fellowship from MICINN (BES-2016-078003).

O.L. and M.S. work was funded by the Spanish Ministry of Science, Innovation and Universities (MCIU/AEI), co-funded by the European Regional Development Fund (ERDF) [SAF2017-82632-P to O.L.]; Autonomous Region of Madrid and co-funded by the European Social Fund and the European Regional Development Fund [Y2018/BIO4747 and P2018/NMT4443 to O.L.]; CNIO is supported by the National Institute of Health Carlos III.

IRB Barcelona and CNIO are Severo Ochoa Award of Excellence recipients from MICINN.

## Author Contributions

N.G-S. designed, performed and analyzed biochemical experiments, except BICD2 pulldowns, done by P.S-F-L. N.G.-S. and P.S-F-L. designed, performed and analyzed cell biology experiments, except mitochondria clustering assays that were performed by J.P. with the contribution of P.S-F-L. L.R. assisted with different cell and biochemical experiments. F.Z. expressed, purified and characterized recombinant BICD2 for structure analysis. M.S. performed further characterization of BICD2, EM experiments and image analysis. O.L. and J.L. provided structural and cell biology expertise, respectively, as well as help designing the study, analyzing data and producing the manuscript. J.R. designed the study, analyzed the data and wrote the manuscript. All the authors reviewed the figures and manuscript.

## Supplementary Figures

**Supplementary Figure S1 (related to Figure 1).**
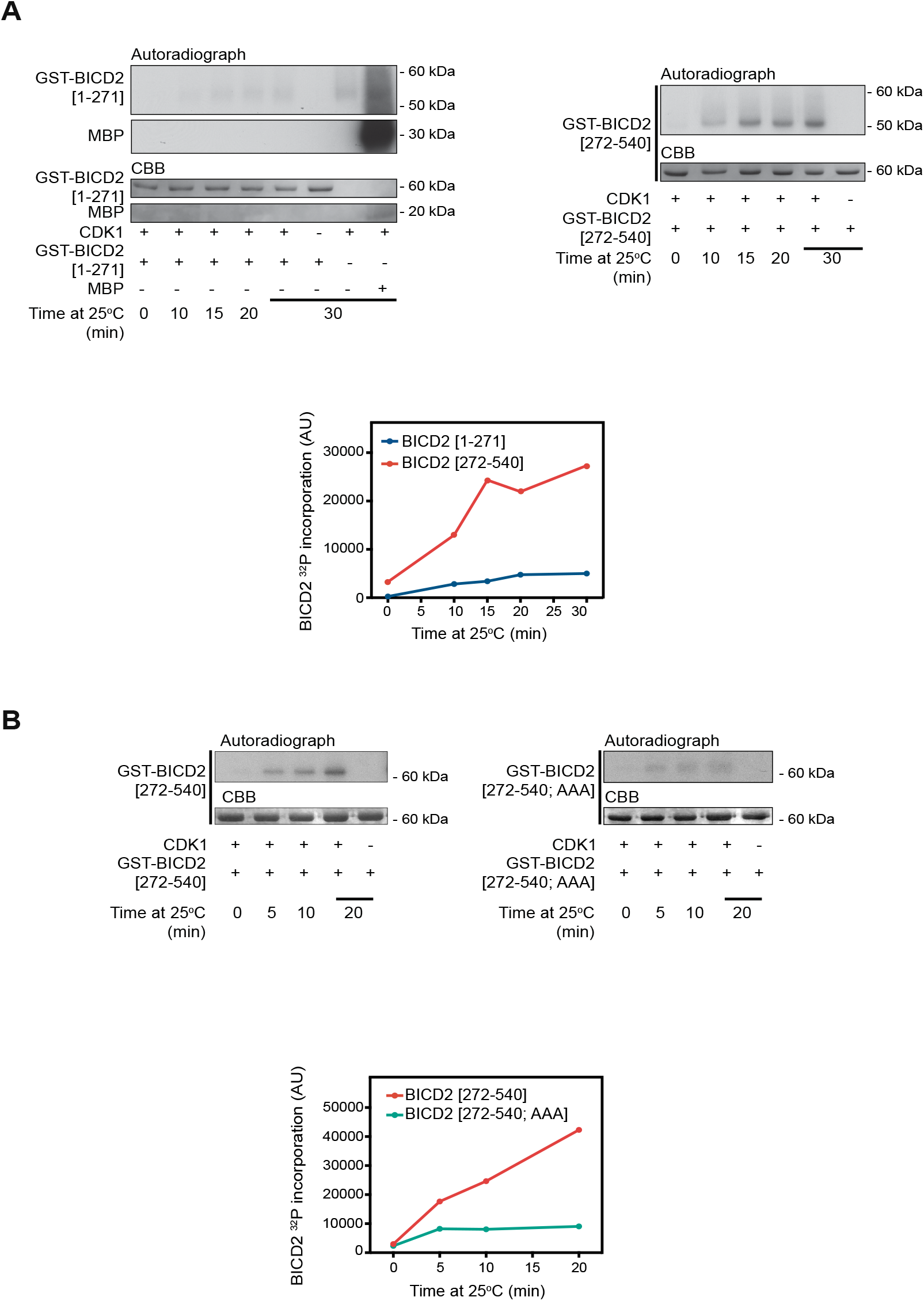
**(A)** *In vitro* phosphorylation of GST-BICD2 fragments by CDK1/cyclin B. Purified bacterial GST-BICD2 [1-271] (left) or GST-BICD2 [272-541] (right) were incubated with recombinant CDK1/cyclin B plus [γ- ^32^P]ATP/Mg^2+^ at 25°C for the indicated times. Myelin basic protein (*MBP*) was used as a positive control. Reactions were stopped with electrophoresis sample buffer and after SDS–PAGE proteins were visualized by Coomassie Brilliant Blue staining (*CBB*) and ^32^P incorporation by autoradiography. Relative ^32^P incorporation was quantified using a Phosphorimager. A non-radioactive replicate of the GST-BICD2 [272-541] reaction at 15 min was used to identify phosphorylated sites by LC-MS/MS. The original analysis using trypsin covered >80% of the sequence and included all the putative CDK1 sites except Thr321, predicted to be contained in a very short peptide (Thr319-Lys324). The analysis identified Ser331 as phosphorylated site. An additional analysis targeted to Thr321 and its surroundings and using chymotrypsin was carried out, detecting an additional phosphopeptide with a single phosphosite ambiguous between Thr319, Ser320 and Thr321. **(B)** Mutation of Thr319, Ser320 and Thr321 strongly interferes with CDK1 phosphorylation of GST-BICD2 [272-541]. GST-BICD2 [272-541] and GST-BICD2 [272-541; T319A, S320A, T321A] (*BICD2 [272-541; AAA]*) were incubated with recombinant CDK1/cyclin B plus [γ-^32^P]ATP/Mg^2+^ at 25°C for the indicated times. Visualization and quantification as in (A).

**Supplementary Figure S2 (related to Figure 1).**
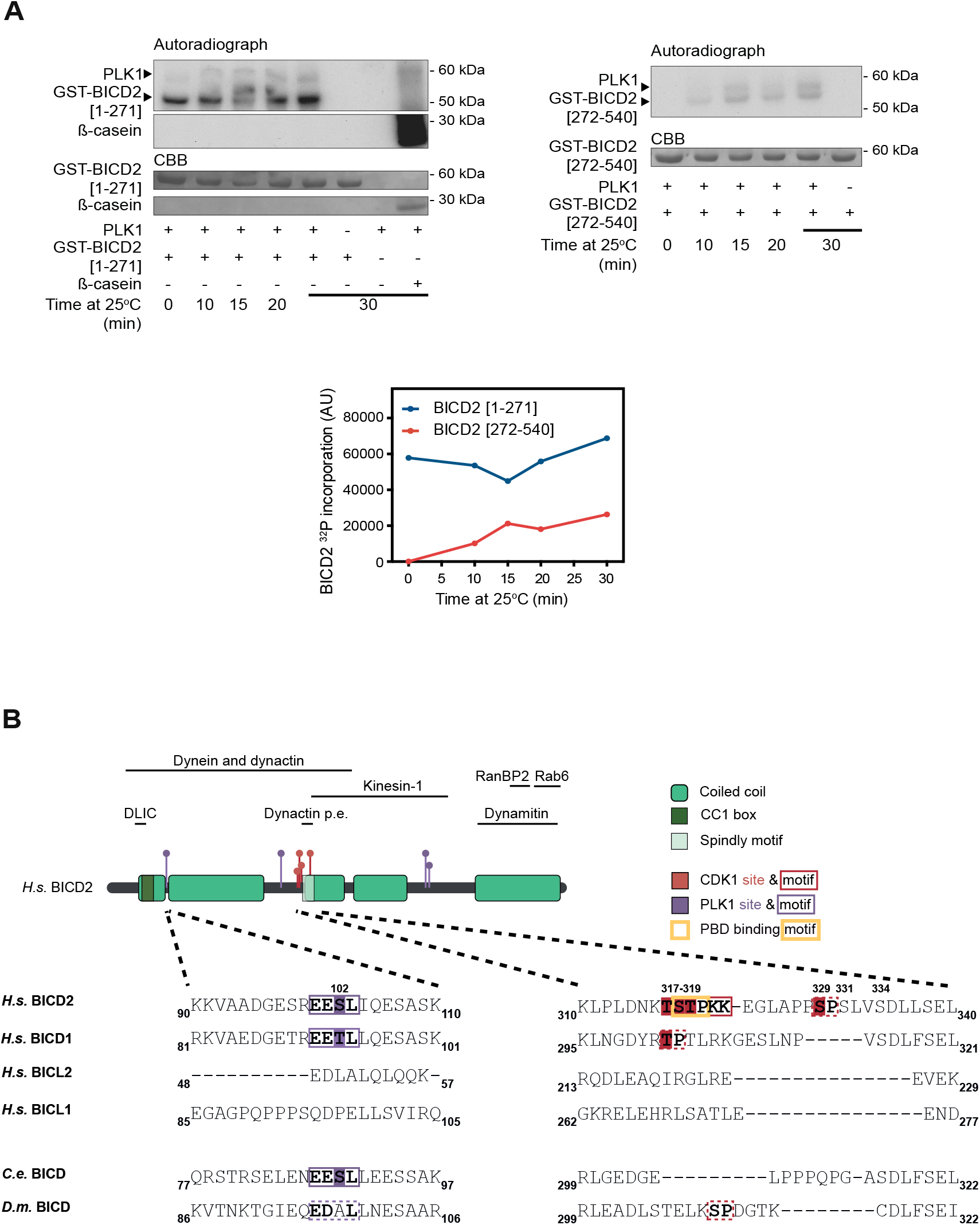
**(A)** *In vitro* phosphorylation of GST-BICD2 fragments by PLK1. Purified bacterial GST-BICD2 [1-271] (left) or GST-BICD2 [272-541] (right), were incubated with recombinant PLK1 plus [γ-^32^P]ATP/Mg^2+^ at 25°C for the indicated times. ß-casein was used as a positive control. Phosphorylation was visualized and quantified as in Supplementary Figure S1A. Note that phosphorylation of GST-BICD2 [1-271] by PLK1 was extremely fast, occurring during the mixing of the reaction components and before it was stopped with electrophoresis sample buffer (t=0). Non-radioactive replicates of the reactions at 15 min were used to identify phosphorylated sites by LC-MS/MS. Analysis using trypsin covered ∼90% of the sequence in both cases. **(B)** *Top,* schematic representation of BICD2, noting regions interacting with different proteins or protein complexes (*Dynactin p.e., dynactin pointed end)*. Coiled coil regions have been predicted using paircoil2 (McDonnell et al., 2006). *Bottom,* sequence alignments of residues surrounding Ser102, Thr317, Ser318, Thr319 and Ser329 in different human (*H.s.*), *Caenorhabditis elegans* (*C.e.*) and *Drosophila melanogaster* (*D.m.*) BICD family members.

**Supplementary Figure S3 (related to Figure 2).**
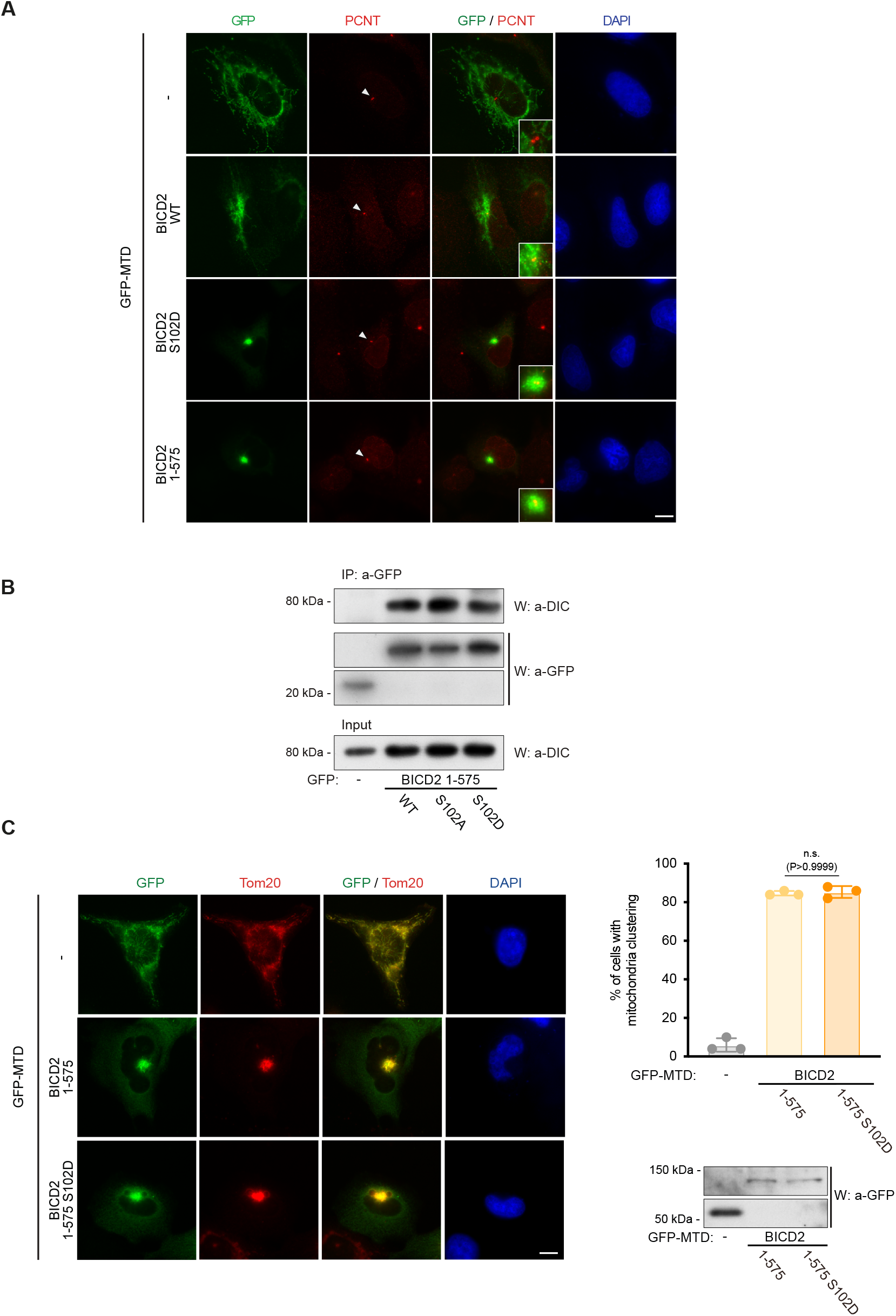
**(A)** Mitochondria-targeting domain (MTD)-fused BICD2 constructs cluster mitochondria around centrosomes. The different GFP- and MTD-tagged constructs used in Figure 2B were transfected in U2OS cells. Immunofluorescent staining was done using anti-GFP and anti-pericentrin (used as a centrosomal marker) antibodies, plus DAPI. Scale bar, 10 µm. **(B)** Phosphonull or phosphomimetic mutations in Ser102 do not affect dynein binding to BICD2 [1-575]. The indicated GFP-fusion proteins were immunoprecipitated from HeLa cells, and analyzed by western blot to detect dynein using anti-dynein intermediate chain (*DIC*), plus GFP. DIC levels in the corresponding extracts are shown in the lower panel. **(C)** A phosphomimetic mutation in BICD2 Ser102 (BICD2 S102D) does not affect dynein mobility towards the centrosomes as detected by mitochondria relocalization and clustering. As in Figure 2B (n=3 biological replicates, 50 cells each; individual replicate means plus mean of replicates ± SD are shown; statistical significance analyzed using a Chi square test). Scale bar, 10 µm. Expression levels of the different polypeptides as detected by western blot are shown.

**Supplementary Figure S4 (related to Figure 4).**
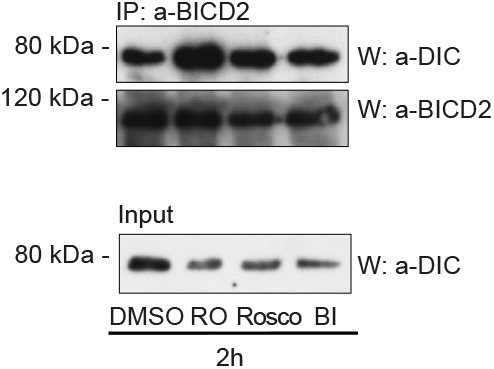
**(A)** Acute CDK or PLK1 inhibition does not significatively disrupts the interaction between BICD2 and dynein in exponentially growing cells. anti-BICD2 immunoprecipitates from exponentially growing HeLa cells treated with the indicated drugs for 2h were analyzed by western blot using anti-DIC or anti-BICD2 antibodies. DIC levels in the corresponding extracts are shown in the lower panel. *RO*, 9 μM RO-3306; *Rosco*, 55 μM roscovitine; *BI*, 100 nM BI2536).

**Supplementary Figure S5 (related to Figure 5).**
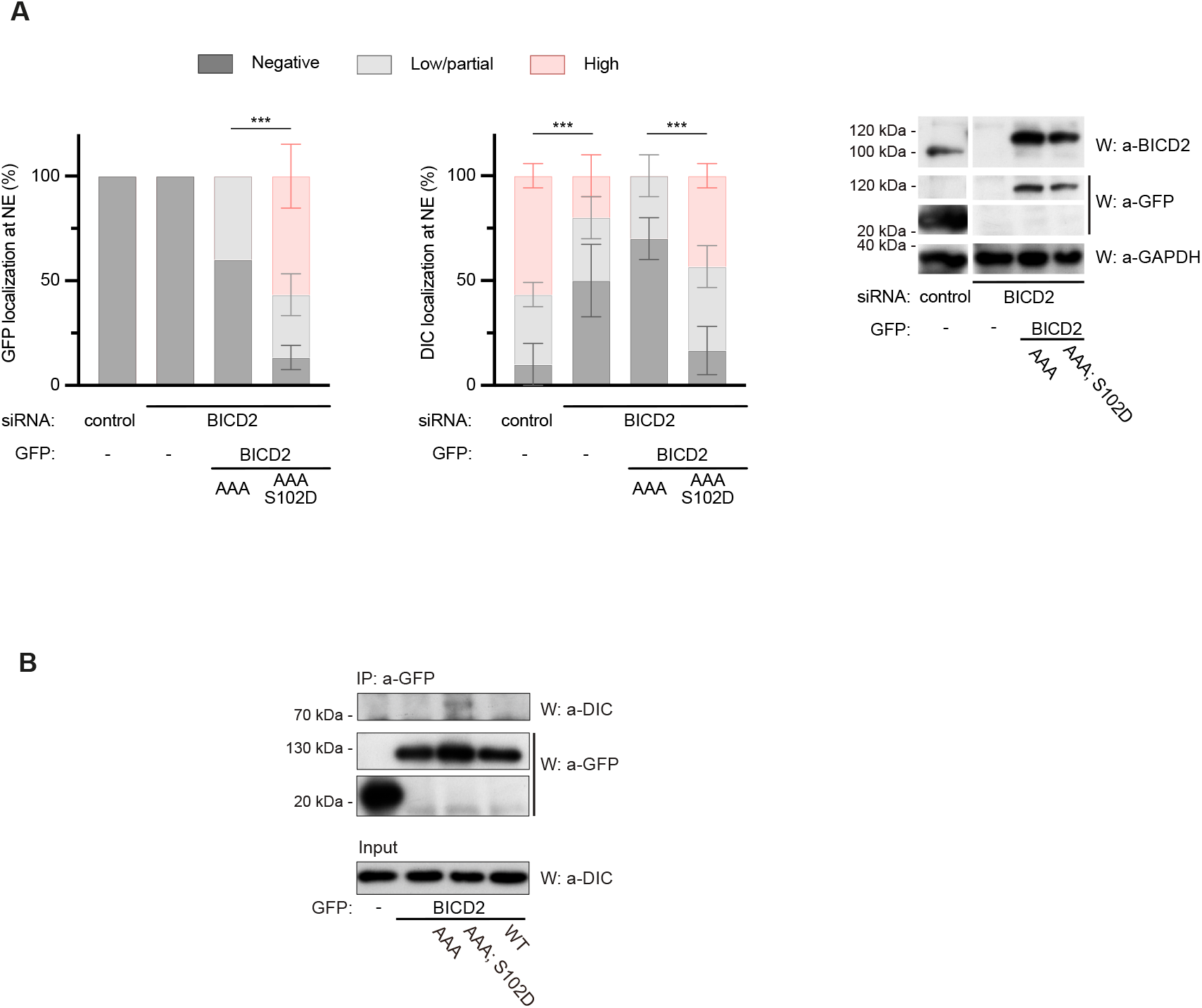
**(A)** A phosphomimetic mutation in BICD2 Ser102 (S102D) is able to rescue dynein nuclear envelope localization in G2 cells with low endogenous BICD2 levels even if major BICD2 CDK1 sites (T319A, S320A, Thr321A) (*BICD2 AAA*) are mutated. HeLa cells were transfected and stained as in Figure 5. GFP-positive G2 cells, identified by positive cyclin B1 cytoplasmic staining, were scored for GFP and DIC at the nuclear envelope and the results quantified as shown (n=3 biological replicates, 10 cells per experiment; mean ± SD is shown; statistical significance was analyzed using a Chi square test). Expression levels of endogenous BICD2 and GFP-fusion proteins are shown. **(B)** The phosphomimetic mutation S102D is able to support dynein binding to BICD2 in a form of the adaptor with mutated BICD2 CDK1 sites (T319A, S320A, T321A). The indicated GFP-fusion proteins were immunoprecipitated from HeLa cells and analyzed by western blot to detect dynein using anti-dynein intermediate chain (DIC) antibodies plus GFP. DIC levels in the corresponding extracts are shown in the lower panel.

